# Spatial enrichment of phosphoinositide metabolism is a molecular switch to promote metastasis

**DOI:** 10.1101/851410

**Authors:** Marisa Nacke, Emma Sandilands, Konstantina Nikolatou, Álvaro Román-Fernández, Susan Mason, Rachana Patel, Sergio Lilla, Tamas Yelland, Laura Galbraith, Lynn McGarry, Jennifer P. Morton, Emma Shanks, Hing Leung, Sara Zanivan, Shehab Ismail, Elke Markert, Karen Blyth, David M. Bryant

## Abstract

The signalling pathways underpinning cell growth and invasion use overlapping components, yet how mutually exclusive cellular responses occur is unclear. We developed 3-Dimensional culture analyses to separately quantify growth and invasion. We identify that alternate variants of IQSEC1, an ARF GTPase Exchange Factor, act as switches to promote invasion over growth by spatially enriching cortical phosphoinositide metabolism. All IQSEC1 variants activate ARF5- and ARF6-dependent PIP5-kinase to promote PI(3,4,5)P_3_-AKT signalling and growth. In contrast, select pro-invasive IQSEC1 variants restrict PI(3,4,5)P_3_ production to discrete cortical domains to form invasion-driving protrusions. Inhibition of IQSEC1 attenuates invasion *in vitro* and metastasis *in vivo*. Induction of pro-invasive IQSEC1 variants and elevated IQSEC1 expression occurs in a number of tumour types and is associated with higher-grade metastatic cancer, activation of PIP3-signalling, and predicts long-term poor outcome across multiple cancers. Spatial enrichment of phosphoinositide metabolism therefore is a switch to induce invasion over growth in response to the same external signal. Targeting IQSEC1 as the central regulator of this switch may represent a therapeutic vulnerability to stop metastasis.

**Highlights:**

- Spatial enrichment of PI(3,4,5)P_3_ is a molecular switch to promote invasion.
- IQSEC1 is a GEF for ARF5/6, promoting PIP5K-dependent PI(3,4,5)P_3_ production downstream of the HGF receptor Met.
- Pro-invasive IQSEC1 variants restrict cortical PI(3,4,5)P_3_ production to subdomains that convert into invasive protrusions.
- IQSEC1 inhibition attenuates *in vitro* invasion and metastasis *in vivo*.
- IQSEC1 module is associated with poor outcome across tumour types.

## Introduction

A central conundrum in biology is that highly overlapping signalling pathways regulate distinct biological outputs. For instance, activation of Receptor Tyrosine Kinases (RTKs), and their downstream effectors such as the MAPK-ERK and PI3K-Akt pathway, can induce either growth or invasion. Since the development of invasive features and the progressive loss of normal tissue organisation are hallmarks of tumour progression ^1^ understanding how cells decode their response to external signals has significant clinical implications.

Metastasis is the major cause of cancer-related death, increasingly recognised as a collective migration event. For a collection of epithelial cells to invade they must undergo rearrangement of normal apical-basal cellular polarisation to clusters or chains of cells lead by an invasive front. The use of 3-Dimensional (3D) culture systems, whereby epithelial cells are embedded in gels of extracellular matrix (ECM) to undergo collective morphogenesis, has illuminated the molecular mechanisms of collective cell polarisation ^2–6^. Such polarity rearrangements can be achieved via altered membrane trafficking of morphogenesis-regulating proteins. For collective invasion to occur, signalling receptors, such as RTKs, must be directed to domains where a pro-invasive ligand is exposed. Some RTKs, such as the HGF receptor Met, require internalisation and endosomal localisation for full oncogenic signalling ^7–9^. Additional membrane trafficking steps, such as recycling back to the cortex, can provide sustained signalling to effectors ^10^. However, what controls whether such signals result in tumour cell growth versus the induction of collective invasion and metastasis remains unclear.

The ARF family of small GTPases (ARF1-6) are implicated in the membrane trafficking and signalling mechanism underpinning single cell invasion ^11–16^. ARF GTPases have fundamental roles in vesicular transport by regulating assembly of coat complexes, lipid-modifying enzymes, and recruiting regulators of other GTPases onto membranes undergoing scission ^17^. ARF6 in particular controls internalisation and recycling of RTKs by acting in concert with a number of GTP Exchange Factors (GEFs) ^18–21^ and GTPase-Activating Proteins (GAPs) ^22,23–26^. Accordingly, small molecule ARF GEF or GAP inhibitors have been used to control invasion *in vitro* and metastasis *in vivo* ^23, 27–29^.

Despite the core requirement for ARF GTPases in membrane trafficking steps controlling RTK signalling, it remains unclear how they could promote an invasion response, rather than growth, from an RTK. Are particular ARF-regulated trafficking pathways induced in invasive cells, such as to enhance endocytic recycling and sustained RTK activation, to promote invasion over growth? Here, we describe that specific pro-invasive transcript variants of the ARF GEF IQSEC1 are upregulated in invasive tumours. These alternate variant proteins act as scaffolds to restrict phosphoinositide metabolism to discrete membrane domains that mature into the invasive protrusion fronts. We identify that IQSEC1 can be targeted to inhibit collective invasion *in vitro* and metastasis *in vivo*.

## Results

### Expression of the ARF GEF IQSEC1 is associated with poor clinical outcome

We examined whether ARF GTPase expression was associated with the acquisition of invasive behaviours. We used 3D culture of prostate cancer cell lines to represent the transition from non-tumorigenic to highly metastatic (Fig 1A). Non-tumorigenic RWPE-1 cells ^30^ formed acini with a central lumen (Fig 1A,B). RWPE-2, an oncogenic *KRAS*-expressing RWPE-1 variant ^30^ formed lumen-lacking aggregates, some of which developed invasive cell chains (Fig 1B, white arrowheads). Bone metastasis-derived PC3 cells ^31^ grew as heterogeneous acini, variably forming round, locally spread, or spindle-shaped invasive cell chains (Fig 1C). Multiday live imaging revealed that invasive chains were derived from single cells that first formed spherical aggregates (growth phase) that initially protruded into matrix, then elongated into chains (Fig 1C, arrowheads, invasion phase).

**Figure 1.**
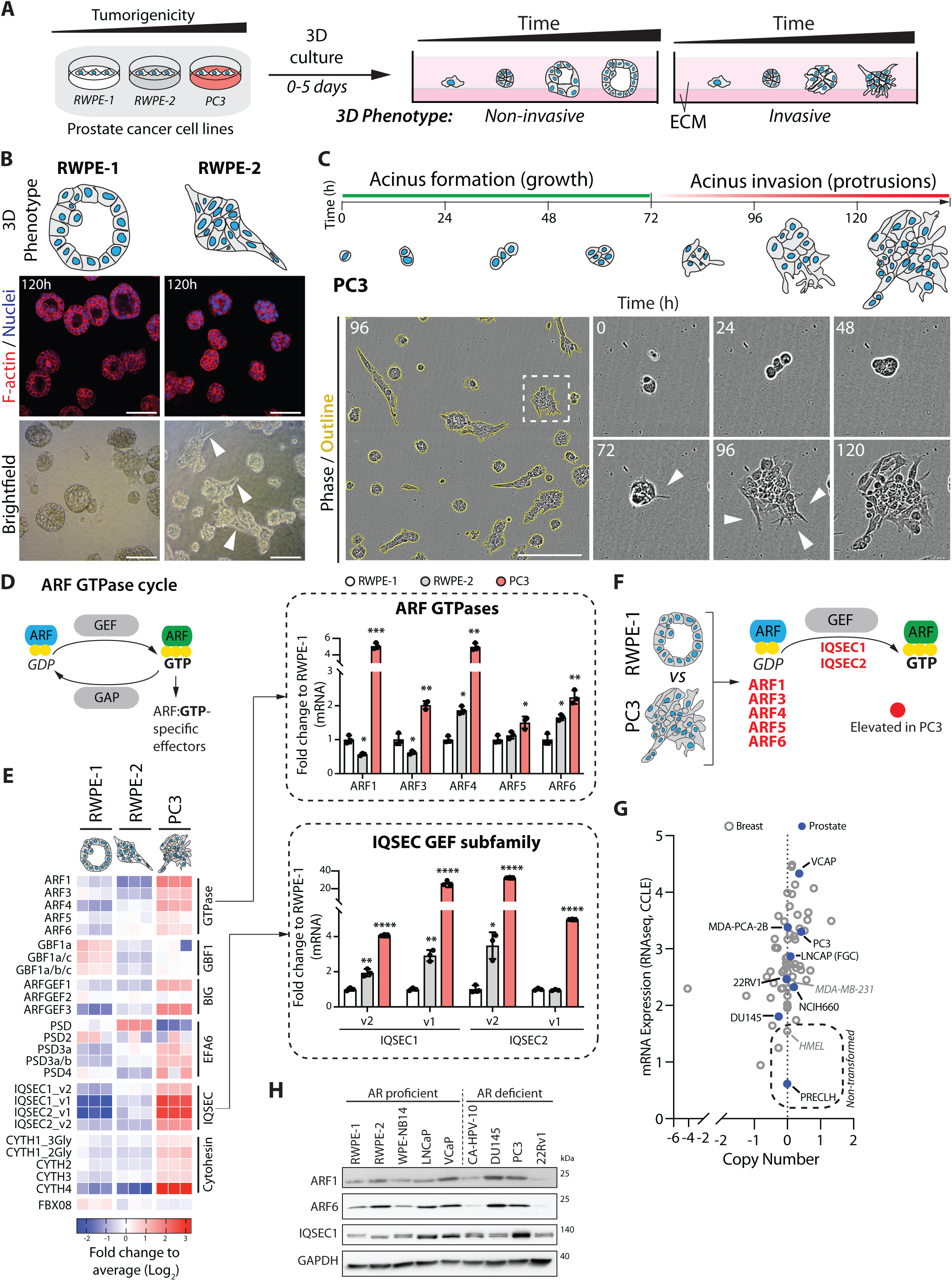
Upregulation of IQSEC1 is associated with tumorigenesis. **(A)** Schema, prostate cell lines forming non-invasive or invasive 3D acini in extracellular matrix (ECM). **(B)** Cartoon, phenotype of typical RWPE-1 and RWPE-2 acini. Confocal (F-actin (red) and nuclei (blue)) and brightfield images show RWPE-1 and RWPE-2 acini at 120 hours. Arrowheads, protrusions. Scale bar, 20μm. **(C)** Schema, PC3 acini form (grow) and invade (protrusions) through ECM over time. Phase contrast images of PC3 acini where higher magnification of boxed region at different time points is shown. Arrowheads, protrusions. Scale bar, 100μm. **(D)** Cartoon, ARF GTPase cycle. **(E)** Heatmap representation of mRNA expression from q-PCR. Data are normalized to RWPE-1 and presented as the log2-transformed fold change compared to the average of all values. Bar graphs summarise fold changes of *ARF* and *IQSEC* mRNA levels. n=3 biological replicates. Values, mean ± s.d. p-values (Student’s t-test). **(F)** Schema, elevated activation of ARF GTPases in PC3 cells by GEFs such as IQSEC1. **(G)** Graph generated using RNAseq data from the Cancer Cell Line Encyclopedia (CCLE) comparing *IQSEC1* gene copy number and mRNA expression levels in multiple breast and prostate cancer and non-transformed cell lines. **(H)** Western blot analysis of androgen receptor (AR) proficient or deficient prostate cell lines using anti-ARF1, ARF6, pan-IQSEC1 isoform and GAPDH (as sample control) antibodies. All p-values: *p≤0.05, **p≤0.01, ***p≤0.001 and ****p≤0.0001.

ARF GTPases function with GEFs and GAPs that control their nucleotide-association state (Fig 1D) ^17^. We observed an association of the GEF *IQSEC1* (also called *BRAG2/GEP100* ^32^) with prostate tumorigenesis. All ARF GTPases, and a number of ARF GEFs were robustly upregulated in PC3 cells (Fig 1E, F). *IQSEC1*, including multiple isoforms, was a robustly induced GEF in RWPE-2 and PC3. Normal prostate and breast cells expressed low levels of *IQSEC1*, while invasive lines (PC3, VCaP) had robustly upregulated *IQSEC1* mRNA, with little variation in copy number (Fig 1G). Western blotting of Androgen Receptor (AR)-proficient and - deficient prostate lines confirmed similar upregulation of ARF6 and IQSEC1 protein expression in metastatic prostate cancer cell lines (LNCaP, VCaP, DU145, PC3) (Fig 1H). *IQSEC2* did not show a similar normal-to-tumorigenic spread across cell lines (not shown). We thus focused on dissecting IQSEC1 molecular function.

### IQSEC1 is a regulator of collective cell invasion

We examined the contribution of IQSEC1 to cell growth and movement. Publicly available *IQSEC1* transcript information revealed multiple variants occurring through combinatorial use of alternate translational initiation sites and alternate splicing (Fig 2A; Table S1). Western blotting suggested simultaneous expression of multiple variants in PC3 cells, with three IQSEC1 bands depleted by IQSEC1 shRNAs (Fig S1A). IQSEC1 depletion reduced proliferation proportional to knockdown efficiency (Fig S1A, B). As PC3 cells grow as a mixed morphology 3D culture (Fig 1C), we developed a machine learning approach to determine whether this heterogeneity also occurred in 2D (Fig S1C). Mirroring 3D collective phenotypes (Fig 1C), single PC3 cells in 2D culture could be classified into round (54%), spread (21%) and spindle phenotypes (17%) (Fig S1D, E). IQSEC1 depletion selectively abolished spindle characteristics, causing increased spread behaviours (Fig S1F).

**Figure 2.**
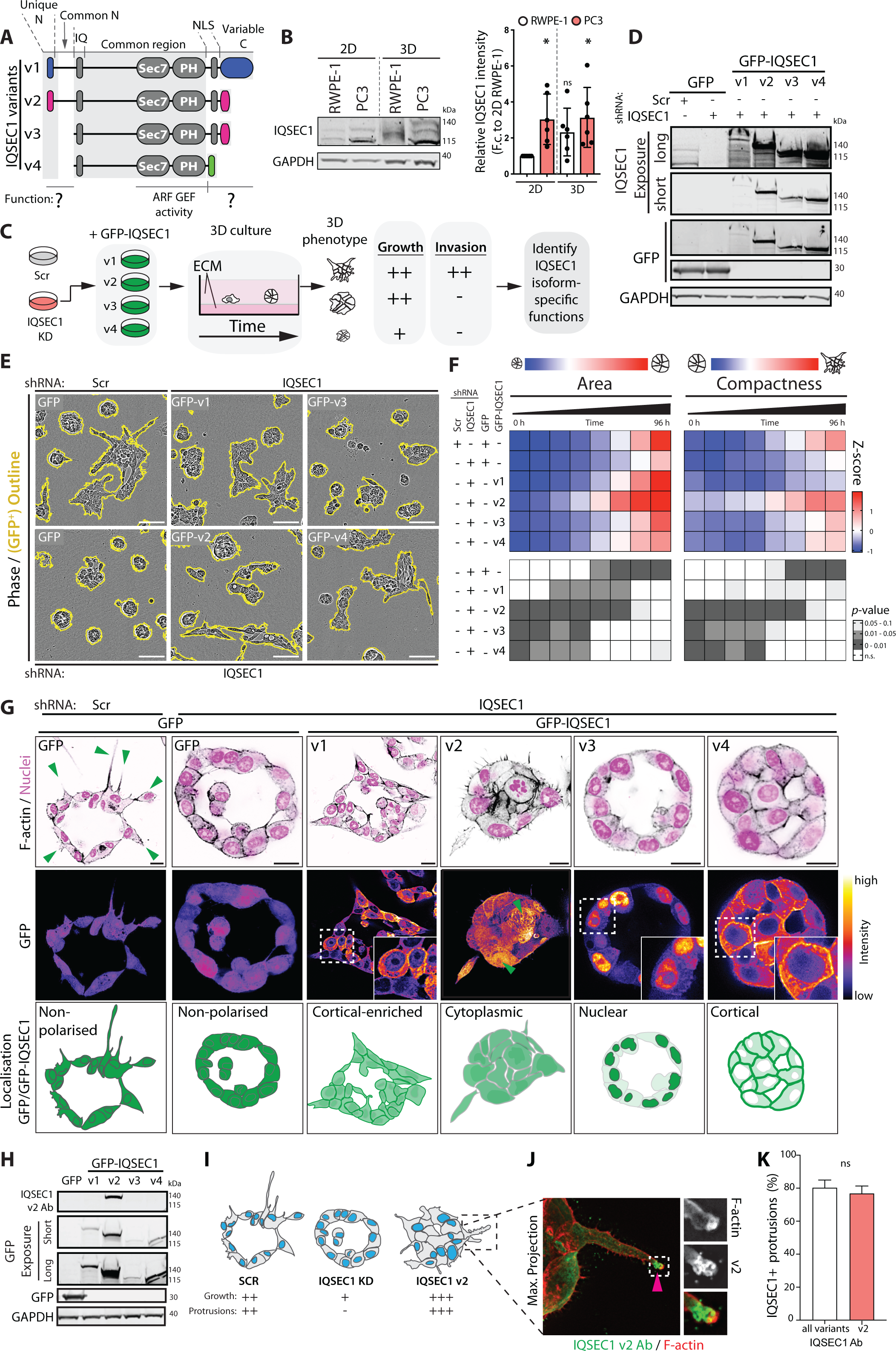
IQSEC1 isoforms differentially regulate collective invasion. **(A)** Schema, domain structure of IQSEC1 variants (v) 1-4. All variants possess an IQ motif, SEC7 and PH domains. v1 and v2 contain N-terminal extensions. A nuclear localization signal (NLS) is present in v1-3, lost in v4. V2/v3 share a common C-terminus. Common domains shown in grey, unique domains indicated by colour; blue; v1, pink; v2 and v3, green; v4. **(B)** Western blot in 2D and 3D using anti-IQSEC1 antibody. The GAPDH blot is is the loading control for the IQSEC1 blot. Relative expression of all IQSEC1 bands normalised to 2D RWPE1 is shown. n=6 independent experiments. Values, mean ± s.d. p-values (Student’s t-test), *p ≤0.05; n.s., not significant. **(C)** Cartoon, isoform specific IQSEC1 functions were identified by comparing the growth and invasion of acini expressing different GFP-IQSEC1 variants upon depletion of endogenous *IQSEC1* (shRNA KD4). **(D)** Western blot of PC3 cells expressing GFP or GFP-IQSEC1 v1-v4, and either Scr or *IQSEC1* KD4 shRNA using anti-IQSEC1, GFP and GAPDH antibodies. All antibodies used on same membrane. Different exposures of IQSEC1 (long, short) are presented to demonstrate expression of all variants. Upper and lower parts of same GFP blot are shown to demonstrate expression of GFP-IQSEC1 variants and GFP control respectively. **(E)** Phase images of acini from cells described in **D**. GFP-positive acini are outlined in yellow. Scale bars, 100μm. **(F)** Quantitation of images shown in **E**. Heatmap shows area and compactness measurements as Z-score-normalised to control (+-+-) values (upper panels). p-values (one-way ANOVA): greyscale values as indicated (lower panels). n=3 independent experiments, 3 replicates/condition, 300 - 650 acini/condition in total. **(G)** PC3 acini described in **D** were stained for F-actin and nuclei (black and magenta respectively in upper panels) after 4 days. Scale bars, 20μm. Localisation of GFP-IQSEC1 can be appreciated from FIRE pseudo-coloured Look Up Table (FIRE LUT) (middle panels, green arrowheads, cytoplasmic localisation). Magnified images of boxed regions are inset. Cartoon, localization of GFP-IQSEC1 variants in PC3 acini (lower panel). **(H)** Western blot of PC3 cells expressing GFP or GFP-IQSEC1 v1-v4 using anti-IQSEC1 (specific for v2) and GAPDH (loading control for IQSEC1 blot) antibodies. Different exposures of GFP (long, short) are presented showing GFP-IQSEC1 proteins. Upper and lower parts of same GFP blot are shown to demonstrate expression of GFP-IQSEC1 variants and GFP control respectively. **(I)** Schema, summary of the effect of *IQSEC1* KD and expression of GFP-IQSEC1 v2 on growth and protrusive ability of PC3 acini. **(J)** Endogenous IQSEC1 v2 co-stained with F-actin (green and red respectively in merge). Magnified images of boxed region are shown. Pink arrowhead indicates localisation at protrusion tip. **(K)** IQSEC1 + protrusion tips were quantified (%/acinus) using antibodies which either detect all variants or are specific for v2. n=2 independent experiments with 49 and 55 acini quantified respectively. Values, mean ± s.e.m. p-values (Student’s t-test): n.s. not significant.

We examined whether IQSEC1-dependent spindle shape was required for cell movement. Live imaging of wounded 2D monolayers revealed cells of various shapes rapidly move into the wound (Fig S1G). IQSEC1-depleted cells displayed a modest defect in migration. In contrast in 3D invasion assays, wounded monolayers embedded in ECM (Matrigel) invade by forming spindle-shaped protrusions that develop into multicellular chains (Fig S1H, white arrowheads). In IQSEC1-depleted cells, although some protrusions formed, multicellular chain formation and invasive activity was strongly compromised (Fig S1H, I). Thus, IQSEC1 is required for growth and multicellular invasive chain formation, resulting in defects in morphogenesis where cell elongation is required (Fig S1J).

### IQSEC1 is a regulator of collective 3D invasion

We dissected IQSEC1 contribution to growth and/or invasion in 3D PC3 acini, which can form without (round) or with (spindle) invasive characteristics (Fig 1C). In order to quantify this growth versus invasive behaviour we developed an automated method of quantitation from hourly imaging of hundreds to thousands of 3D acini over multiple days (see Materials and Methods and Fig S2A).

Elevated IQSEC1 expression in PC3 cells, compared to RWPE-1 cells, was maintained upon plating of both cell types into 3D culture (Fig 2B). Growth of acini from single cells could be measured by increased area over time (Fig S2B, C). The progressive development of protrusive invasion could be determined using a ‘Compactness’ measurement (Fig S2B, C, S1 and S2 Movies). Similar to 2D, IQSEC1 depletion modestly decreased 3D acinar growth (Fig S2C), proportional to depletion levels (Fig S1A). Mirroring 3D wound invasion defects (Fig S1H), the most prominent effect of IQSEC1 depletion was abolished protrusive invasion (Fig S2B, C).

We examined whether protrusion-forming activity was a common feature of all IQSEC1 isoforms. We focused on the four annotated variants (v1-4; Fig 2A). IQSEC1 v1-4 possess the IQSEC family-defining features of a calmodulin-binding IQ domain, a catalytic SEC7 ARF GEF domain, and a lipid-binding PH domain. Alternate initiation sites provide v1 or v2 with unique N-termini, followed by a common N-terminal extension. Three alternate C-termini can occur, with the v4-type tail truncating a potential nuclear localisation sequence.

We stably, individually restored each RNAi-resistant GFP-tagged IQSEC1 variant in the background of depletion of all endogenous IQSEC1 transcripts. All IQSEC1 variants could restore growth and/or invasion to levels matching or above control cells (Fig 2C-F). Variants containing the N-terminal extension (v1, v2) conferred the strongest effect on growth. In contrast, protrusive activity occurred in an isoform-selective manner: only v2 restored spindle behaviours in both 2D and 3D to levels surpassing controls (Fig 2E, F, S2D). To corroborate the live imaging approach we examined fixed acini through 3D confocal imaging. Restoration of v2 to IQSEC1 KD cells increased the total number of nuclei, the level of a proliferation marker (Ki67) and suppressed apoptosis without changing total acinus volume (Fig S2E-H). This represented formation of protrusions and disruption to lumen formation. In contrast, v4 failed to rescue IQSEC1 KD-induced growth defects, but instead increased area by increasing acinus volume without changing cell number, due to the presence of a lumen. Thus, IQSEC1 v2 (also known as BRAG2b) is a major regulator of 3D growth and invasion.

We explored the IQSEC1 v2 properties that promote invasion. IQSEC1 can function in ARF GTPase-dependent endocytosis at the cell cortex, and in the nucleus to control nucleolar architecture ^32, 33^. V3 showed predominantly nuclear localisation, while v4 was cortical (Fig 2G, S2I). V1 showed mixed cortical and cytoplasmic localisation. In contrast, v2 displayed a mix of cytoplasmic, cortical and nuclear labelling (Fig 2G, green arrowheads, S2I). We reasoned that in overexpressed GFP-tagged variants may mask vesicular pools. Accordingly, anti-IQSEC1 antibodies directed to either all isoforms or to v2-specifically (Fig 2H) labelled both the cytoplasm and tubulovesicular compartments behind the tips of invasive acinar protrusions in 3D (Fig 2I-K, arrowheads, S2J, K). Thus, one locale of IQSEC1 v2 function is focally at endosomes in protrusion tips (Fig 2I-K).

We mapped the domains of v2 responsible for its localisation and potent invasion-inducing activity. We expected that this would be conferred by the unique v2 N-terminus. However, using mutants and chimeras of IQSEC1 isoforms revealed a nuanced and combinatorial effect of alternate N- and C-termini on IQSEC1 localisation and protrusion induction (Fig S3A-G). Surprisingly, the unique v2 N-terminus (2N) was not required for invasion. Any variant containing the v2-tail and *any* N-terminal extension promoted invasion. Accordingly, replacing the unique v2 N-terminus (2N) with that from v1 (1N) even enhanced invasion over and above that of v2 (Fig S3E-G). This was concomitant with a shift away from nuclear or cortical localisation, towards the cytoplasm (Fig S3C, 1N/2C chimera). This is in contrast to expression of v1 which did not induce invasion (Fig 2F). This is due to the presence of a v1-type C-terminus (1C) which in all experiments inhibits invasion, concomitant with a more general cortical recruitment. A v2 tail without N-terminal extension (i.e. v3) similarly does not promote invasion. This maps the invasion-inducing activity of IQSEC1 to the N-terminal extension that is common to only v1 and v2. However, the type of C-terminus influences whether this N-terminus induces invasion (Fig S3A-G). Mutational inactivation of GEF activity (IQSEC1 v2^GEF*^) confirmed that IQSEC1 v2-induced spindle shape in 2D (Fig S3B) and protrusive invasion in 3D (Fig S3D-G) are due to its ability to activate ARF GTPases. Thus, the N-terminal common extension in v2 confers enhanced invasive activity in a GEF-dependent manner.

### IQSEC1 activates ARF5/6 in distinct locations within protrusions

IQSEC1 functions with both ARF5 and ARF6 to control integrin endocytosis and focal adhesion disassembly^13, 34, 35^. Given IQSEC1 GEF activity-dependency for invasion (Fig S3D-G), we examined whether ARF5 and ARF6 are required for this process. While individual ARF depletion caused some changes that mimicked IQSEC1 loss, only combined ARF5/6 depletion reduced all aspects of 2D spindle shape and 3D growth and invasion (Fig 3A-C, S4A). We examined whether individual or co-overexpression (OX) of ARF5 or ARF6, wild-type (WT) or fast-cycling mutants (Fig S4B, C) ^13, 36^ induced invasive behaviours. Only ARF5/6 WT co-overexpression increased 2D spindle shape, 3D growth and 3D invasion (Fig 3D-F, S4D-G). While fast-cycling mutants induced some 2D spindle behaviours, they failed to induce 3D effects, either alone or in combination, indicating that normal GTPase cycling was required. Crucially, IQSEC1 depletion abolished all effects of ARF5/6 co-overexpression (Fig 3D-F, S4H). This identifies IQSEC1 as the major GEF regulating growth and invasion with ARF5/6 as its major targets.

**Figure 3.**
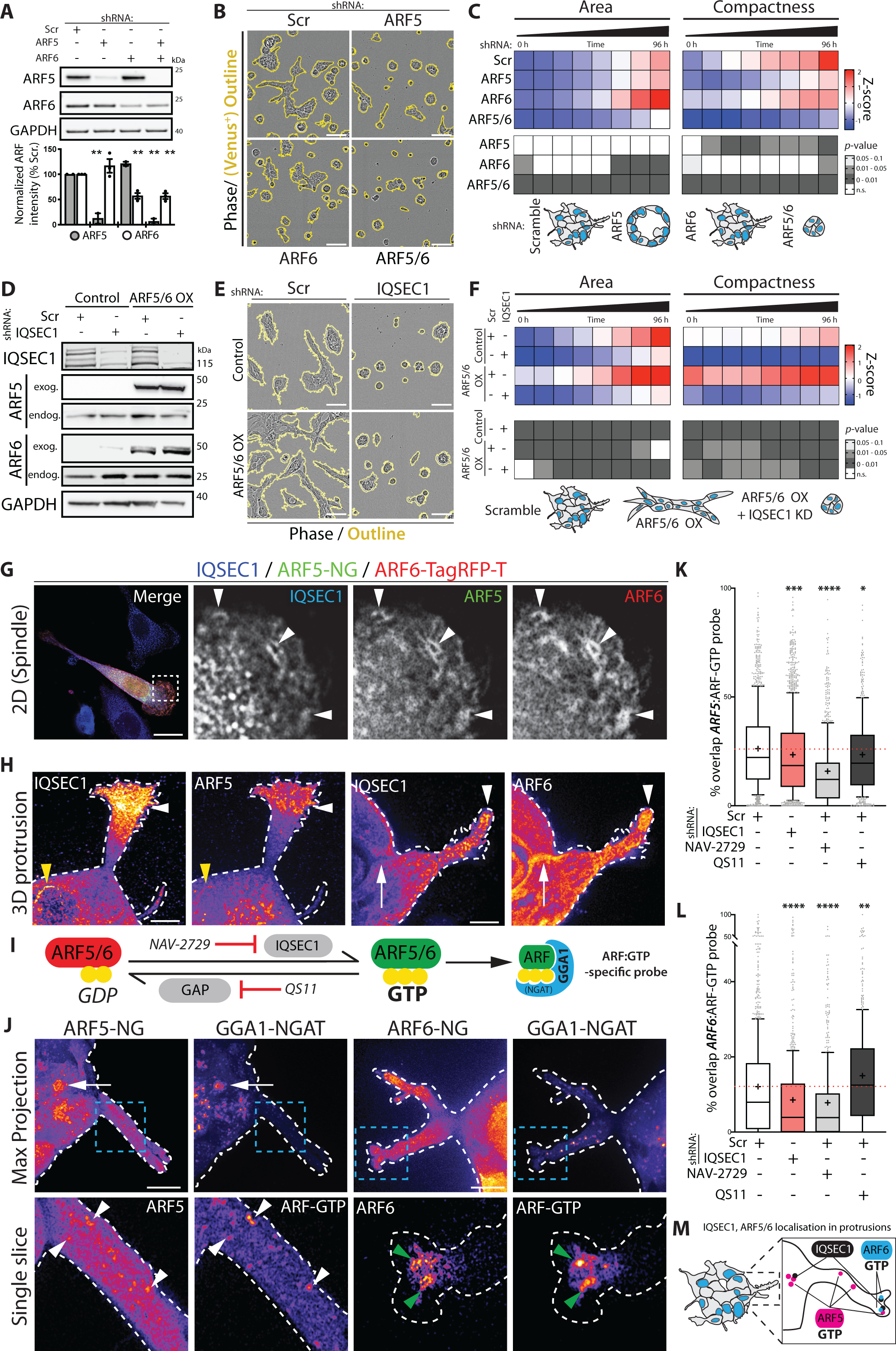
IQSEC1 activates ARF5/6 in distinct locations within protrusions. **(A)** Western blot of PC3 cells expressing Scr, *ARF5* or *ARF6* shRNA using anti ARF5, ARF6 and GAPDH (loading control for ARF6 blot) antibodies. ARF intensity normalised to Scr is quantified. n=3 independent experiments. Values, mean ± s.d. p-values (one-way ANOVA): **p≤0.01. **(B)** Phase images of PC3 acini described in **A**. Yellow outlines indicate shRNA (mVenus) positive acini. Scale bars, 100μm. **(C)** Quantitation (of images in **B**) is shown for area and compactness measurements as Z-score-normalised values (upper heatmap). p-values (one-way ANOVA): greyscale values as indicated (lower heatmap). n=3 independent experiments, 5 replicates/condition, 2,880 – 3,188 acini quantified/condition in total. Cartoon, acini phenotype representative of each condition. **(D)** Western blot of PC3 cells co-overexpressing (OX) mNeonGreen (mNG) and TagRFP-T (RFP) (Control) or ARF5-mNG and ARF6-RFP (ARF5/6), and either Scr or *IQSEC1* KD4 shRNA. Anti-IQSEC1, ARF5, ARF6 and GAPDH (loading control for ARF5 blot) antibodies were used. Both endogenous and exogenous (OX) ARFs were detected. **(E)** Phase images of PC3 acini described in **D**. Scale bars, 100μm. **(F)** Quantitation of images shown in **E**. n=2 independent experiments, 4 replicates/condition, 1,254 – 1,567 acini/condition in total. **(G)** PC3 cells co-overexpressing ARF5-mNG and ARF6-RFP (ARF5/6 OX) were stained for IQSEC1 v2 and merged image of spindle shaped cell shown (left panel). Magnified images of boxed region are shown. White arrowheads indicate areas of colocalization. Scale bars, 20μm. **(H)** PC3 acini expressing either ARF5-mNG or ARF6-mNG were stained for IQSEC1. Localisation of these proteins is shown using FIRE LUT. Yellow and white arrowheads, colocalization in juxtanuclear region and protrusive tips, respectively. White arrows, lack of colocalization. Scale bars, 5μm. **(I)** Schema, GTPase cycle of ARF5 and ARF6, site of action of the IQSEC1-inhibiting molecules NAV-2729, the pan ARFGAP-inhibitor QS11, and GTP-loaded ARF detection by a GGA1-NGAT probe. **(J)** PC3 acini expressing either ARF5-mNG or ARF6-mNG and GGA1-NGAT-RFP were fixed and FIRE LUT of maximum projections (upper panels) and a single Z-slice of boxed regions (lower panels) shown. White and green arrowheads, colocalization in protrusions, white arrows, colocalization in cell body. Scale bars, 10μm. **(K-L)** PC3 cells expressing **(K)** ARF5-NG and GGA1-NGAT or **(L)** ARF6-NG and GGA1-NGAT were transfected with either Scr or *IQSEC1* KD4 shRNA. Cells were treated with NAV-2729 or QS11 for 24 hours. Quantitation of % overlap of ARF and ARF-GTP probe per cell is shown in box-and-whiskers plot: 10–90 percentile; +, mean; dots, outliers; midline, median; boundaries, quartiles. n=2 independent experiments, 3 replicates/condition with a minimum of 500 cells/condition in total. p-values (one-way ANOVA): *p≤0.05, **p≤0.01, ***p≤0.001 and ****p≤0.0001. **(M)** Schema, localisation of IQSEC1 and active ARFs in protrusions.

Despite identifying IQSEC1 GEF activity as essential for growth and invasion (Fig S3E-G) we did not observe global defects in GTP-loading of ARF5 or ARF6 upon IQSEC1 depletion (Fig S4I). As IQSEC1 v2 localised to a discrete pool at tips of protrusions (Fig 2J), we reasoned that IQSEC1 may regulate a small but crucial ARF5/6 pool. We developed an imaging-based approach to determine the localization of the IQSEC1-ARF complex. Endogenous IQSEC1 v2 localised with ARF5-mNeonGreen and ARF6-TagRFPT in tubulovesicular structures near the leading edge lamellipodium of spindle 2D cells (Fig 3G, white arrowheads). In 3D, ARF5 displayed a prominent vesicular localisation in the acinus body, which overlapped with IQSEC1 in some regions, while ARF6 was prominent at cell-cell contacts lacking IQSEC1 (Fig 3H, yellow arrowheads and white arrows respectively). Both ARF5 and ARF6 could be found with IQSEC1 v2 in protrusion tips (Fig 3H, white arrowheads).

To identify the location of active ARF we expressed a sensor that detects GTP-loaded ARF proteins (GGA1 NGAT domain ^37^) (Fig 3I). GGA1 NGAT-TagRFP-T colocalized with ARF6 most notably in the tips of protrusions (Fig 3J, green arrowheads), and with ARF5 in puncta in the body of the protrusion (white arrowheads) and the acinus body itself (white arrows).

To confirm that IQSEC1 was controlling this localized GTP loading of ARF5/6 we developed an automated method to analyse the percentage of overlap of the GGA1-NGAT probe with ARF5- or ARF6-positive puncta. We validated the sensitivity of this approach by first examining whether chemical inactivation of ARF GEF or GAP activity could be detected (Fig 3I). We identified that a reported ARF6 inhibitor (NAV-2729) ^29^, but not the Cytohesin family ARF GEF inhibitor (Secin-H3) ^38, 39^, also reduces IQSEC1-catalysed nucleotide exchange on ARF5 (Fig S4J, K). In cells, NAV-2729 significantly decreased GGA1-NGAT recruitment to both ARF5 and ARF6-positive puncta (Fig 3K, L). In contrast, the ARF GAP inhibitor QS11 significantly increased GGA1-NGAT recruitment to ARF6, while decreasing recruitment to ARF5. Thus, our approach sensitively detects GTP-loading of specific ARF subpopulations (Fig 3J-L). Applying this approach in IQSEC1-depleted cells we observed a robust decrease, but not complete loss, of recruitment of GGA1-NGAT to both ARF5 and ARF6 (Fig 3J-L). These data confirmed that IQSEC1 controls GTP loading of a pool of ARF5/6 that is crucial for invasive activity (Fig 3M).

We determined the morphogenetic effect of modulating ARF activity. Secin-H3 treatment, which minimally acts on IQSEC1-mediated ARF activation (Fig S4K) modestly decreased 3D growth, without affecting invasion (Fig S4L-O). ARFGAP inhibition (QS11) ^27, 39^ increased 3D invasion, without affecting growth. However, in line with being a dual IQSEC1-ARF5/6 inhibitor, NAV-2729 treatment abolished 3D growth and invasion. Thus, ARF5 and ARF6 are the major targets of IQSEC1 GEF activity. Chemical or genetic inhibition of IQSEC1-directed ARF activation strongly attenuates 3D growth and invasion.

### IQSEC1 is a scaffold for Met and Akt signalling

All IQSEC1 variants possess the SEC7-PH domains required to activate ARF GTP loading (Fig 4A). The enhanced invasion-promoting capabilities of IQSEC1 v2 are conferred by the shared N-terminal extension (encoded by Exon 3; Fig 4A). We used mass spectrometry protein-protein interaction analysis of IQSEC1 chimeras to identify domain-specific IQSEC1 binding partners. We classified interactors as unique to v1 or v2 N-termini (1N or 2N), shared between v1 or v2 N-terminal extension (Exon 3), to the core IQ-SEC7-PH region (Core), or specific to the v2 C-terminus (2C) (Fig 4A). We prioritised interactors where multiple components of a complex could be identified. We ordered these by mRNA expression change from two pairs of PC3-derived sublines which possess epithelioid (PC3E, Epi) compared to mesenchymal properties (GS689.Li, EMT14) ^40,41^, to focus on invasion regulators (Fig 4A-C). Using this approach, we identified the interaction of IQSEC1 with a number of oncogenic signalling pathways (mTORC2, RAF, NFkB), cytoskeletal and scaffolding complexes (Utrophin, 14-3-3), a phosphatase (PP6), and trafficking complexes (EARP, AP3/ARFGAP1). In addition to the known interactors Calmodulin-1/-2, we also identified three transmembrane receptors (Met, LRP1 and SORL1), and other factors (TUBB6, FHOD1). We confirmed a number of novel IQSEC1 interactors in independent immunoprecipitations, and interactor protein level alterations by western blotting of a series of PC3-derived sublines with varying invasive abilities (Fig 4D, S5A-C).

**Figure 4.**
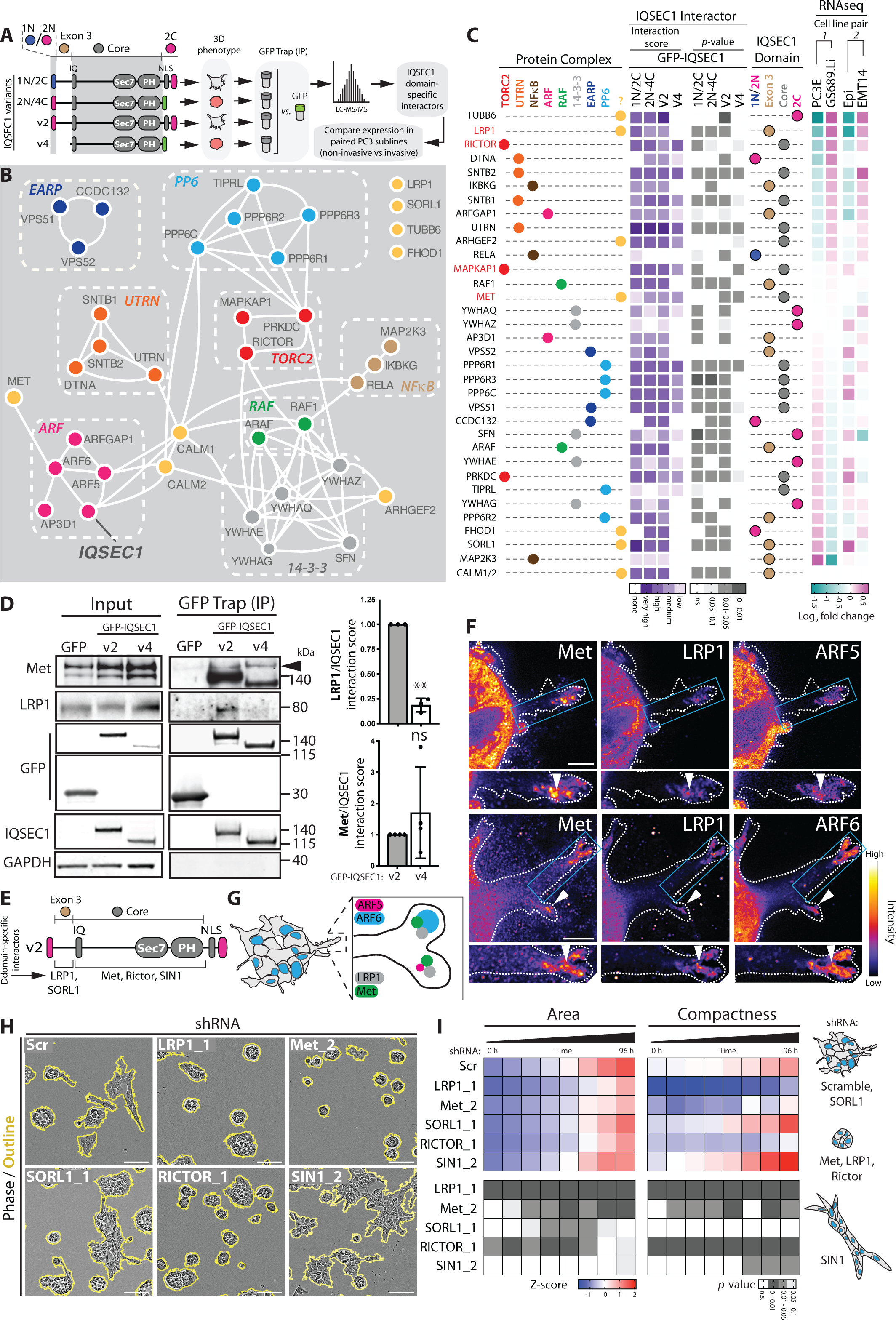
IQSEC1 is a scaffold for Met signalling. **(A)** Schematic, IQSEC1 chimeras and variants display distinct phenotypes in 3D culture. GFP-trap immunoprecipitation was performed on PC3 cells expressing these proteins followed by tryptic digestion “on-beads” and LC/MS/MS analysis. IQSEC1 domain-specific interactions were identified and sorted by mRNA expression compared in paired PC3 subclones. **(B)** STRING network analysis of IQSEC1 binding partners identified by MS visualised using Cytoscape. Most IQSEC1 binding partners could be clustered into 8 protein complexes (colour coded). **(C)** Schema, indicates the protein complex each binding partner is associated with. Heatmap shows the fold change of IQSEC1 interactors from panel **B** binding to different IQSEC1 domains over GFP control. Values are −log_2_(FC), very high = −7 to −5, high = −5 to −2.5, medium −2.5 to −1, low = −1 to −0.05, no binding >0.05. p-values are indicated in grey heatmap. The specific IQSEC1 domain to which each interactor binds is also depicted. Interactions were sorted according to the fold change in mRNA of non-invasive PC3 subclones (GS689.Li, EMT) compared to invasive subclones (PC3E, Epi) (RNAseq). **(D)** GFP-trap immunoprecipitation was performed on PC3 cells expressing GFP, GFP-IQSEC1 v2 or v4. Western blot analysis was then carried out using anti-Met, LRP1, GFP, IQSEC1 and GAPDH antibodies. GAPDH is the loading control for the LRP1 blot. Quantitation of LRP1/IQSEC1 and Met/IQSEC1 interactions for each GFP-trap are shown. n=3 and n=4 independent experiments, respectively. Values, mean ± s.d. p values (one-way ANOVA): **p≤0.01 and n.s. not significant. **(E)** Cartoon, depicts domain specific binding of IQSEC1 interactors. **(F)** PC3 acini expressing ARF5-mNG or ARF6-mNG were stained for Met and LRP1. FIRE LUT images are displayed with magnified images of boxed regions shown in lower panels. White arrowheads indicate localisation in protrusion tips. Scale bars, 5µm (upper) and 10µm (lower). **(G)** Schema, colocalization of IQSEC1-ARF interactors in protrusive tips. **(H)** Phase images of PC3 acini expressing Scr, *LRP1_1, Met_2, SORL1_1, RICTOR_1* or *SIN1_2* shRNA after 96 hours. Scale bars, 100μm. **(I)** Quantitation of images shown in H. Heatmaps show area and compactness measurements as Z-score-normalised values (upper panels). p-values (one-way ANOVA): greyscale values as indicated (lower heatmaps). n=2 independent experiments, 4 replicates/condition, 700 - 2,400 acini/condition in total. Cartoon, depicts acinus phenotype representative of each condition.

We initially focused on transmembrane interactors of IQSEC1. IQSEC1 associated with Met, the receptor for HGF. This occurred through the core region, indicating that all IQSEC1 variants can potentially interact with Met (Fig 4D, E, S5A). In contrast, two proteins interacted only with the invasion-inducing regions of IQSEC1: LRP1 (Low density lipoprotein receptor-Related Protein 1) and SORL1 (Sortilin-Related Receptor 1), members of the LDL Receptor family (Fig 4D, E, S5A). LRP1 and SORL1 protein, however, displayed a mutually exclusive expression: LRP1 expression was elevated in invasive PC3 sublines (EMT14, GS689.Li), whereas SORL1 was associated with epithelial sublines (PC3E, Epi) and was anti-correlated with active pY1234/1235-Met (pMet) levels in cells (Fig S5B, C). In addition, LRP1 and Met both localised to the tips of invasive protrusions in 3D (Fig 4F, G), and formed a complex irrespective of IQSEC1 expression levels (Fig S5D). These data suggest that LRP1 and Met form a complex in the tips of protrusions onto which IQSEC1 associates to promote invasion.

Supporting the enrichment of active Met and LRP1 in invasive PC3 sublines (Fig S5A-C), depletion of LRP1 or Met reduced spindle shape in 2D and attenuated 3D growth and invasion (Fig 4H, I, S5E-G). LRP1 is a 600kDa protein that is cleaved to release a 515kDa extracellular fragment and an 85kDa transmembrane receptor ^42, 43^. Expression of a GFP-tagged transmembrane LRP1 fragment (LRP1^TM^-GFP) was not sufficient to induce invasion, suggesting both transmembrane and extracellular LRP1 fragments are required for invasion (Fig S5H). In contrast, SORL1 depletion increased spindle behaviours in 2D but did not significantly affect invasion or growth in 3D (Fig 4H-I, S5G, I-K). These data support a positive role for LRP1/Met in invasion, while SORL1 may be antagonistic.

We examined downstream signalling from Met that drives invasion. Although we identified RAF protein association with IQSEC1, MEK-ERK inhibitors minimally affected PC3 cell invasion (not shown). We thus focused on signalling to the TORC2-Akt pathway, as multiple subunits of this complex (RICTOR, SIN1/MAPKAP1, PRKDC) associated with IQSEC1. RICTOR is an essential positive regulator of mTORC2 ^44, 45^, while SIN1 has been reported to inhibit mTORC2 activity ^46^. Accordingly, RICTOR depletion robustly abolished Akt S473 phosphorylation (pAkt), spindle shape in 2D, and 3D invasion and growth (Fig 4H, I, S5G, J-L). Notably, while SIN1 depletion decreased spindle behaviours in 2D, it robustly increased invasion in 3D, suggesting 3D-specific differences in this pathway (Fig 4H, I, S5G, M). Finally, we examined ARFGAP1 as a potential antagonist of IQSEC1-ARF function. Unexpectedly, ARFGAP1 depletion phenocopied IQSEC1 depletion: decreasing pAkt levels, spindle shape in 2D, modestly decreasing 3D growth, and profoundly inhibiting 3D invasion (Fig S5G, N-P). This suggests ARFGAP1 as an effector of ARF5/6 in regulating invasion and signalling to the mTORC2-Akt pathway (Fig S5Q), similar to observation of dual effector-terminator functions of other ARFGAPs ^47–49^. Thus, an IQSEC1-ARF5/6-ARFGAP1 complex may control LRP1-Met signalling to mTORC2 during formation of invasive 3D protrusions.

### An IQSEC1-LRP1 complex modestly regulates Met endocytic trafficking

Oncogenic Met signalling requires internalization^9^. We investigated whether IQSEC1 controls HGF-Met signalling to Akt by controlling Met endocytic trafficking. HGF stimulation increased 3D growth and robustly induced invasion, which was substantially blunted by IQSEC1 depletion (Fig S6A-D). ARF5/6 co-overexpression resulted in a modest but consistent increase in pAkt levels, which was nonetheless attenuated to levels similar to parental cells by IQSEC1 depletion (Fig 3D, 5A,B, S6A, B). This suggests that IQSEC1-ARF5/6 control signalling to the Akt pathway.

**Figure 5.**
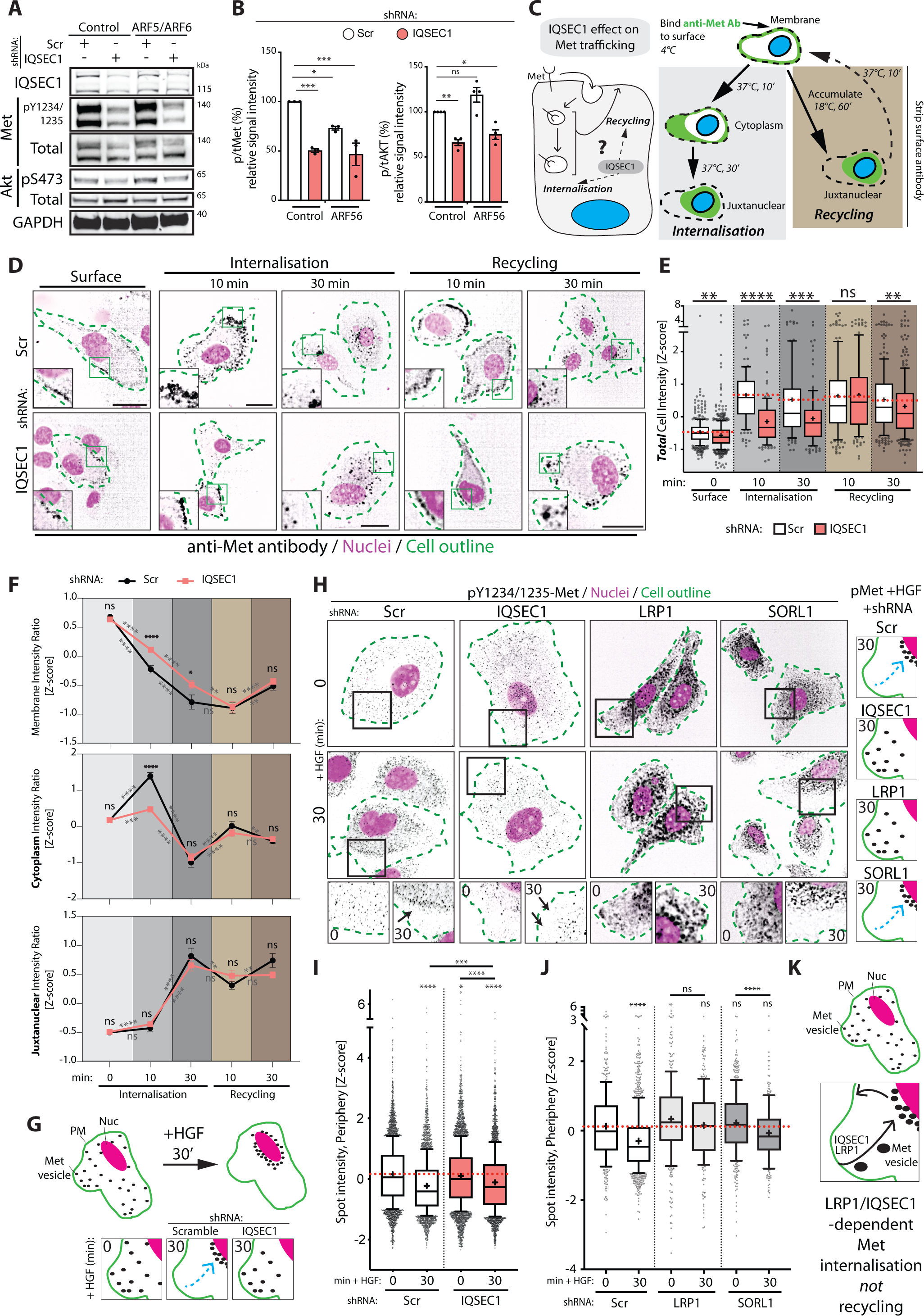
IQSEC1-LRP1 complex regulates Met endocytic trafficking. **(A)** Western blot of PC3 cells co-expressing mNG and RFP (Control) or ARF5-mNG and ARF6-RFP (ARF5/6) with either Scr or *IQSEC1* KD4 shRNA. Anti-IQSEC1, phospho-Y1234/1235 Met, Met, phospho-S473 Akt, Akt, and GAPDH (sample control) antibodies were used. **(B)** Quantitation of phospho/total Met and phospho/total Akt expression is presented as signal intensity relative to control. n= 3 independent experiments. Values, mean ± s.d. p values (one-way ANOVA). **(C)** Schema, effect of IQSEC1 on Met trafficking. **(D)** PC3 cells expressing Scr or *IQSEC1 KD4* shRNA were incubated with a Met-647 fluorescent antibody at 4°C (Surface) prior to stimulation with HGF (Internalisation). Cells were treated with chloroquine to allow accumulation of surface-derived Met (black) for 1 hour at 17°C prior to HGF stimulation (Recycling). Cells were stained with F-actin (green outlines) and Hoechst (nuclei in magenta). Magnified images of boxed regions are inset. Scale bars 20µm. **(E)** Intensity of Met in each cell was quantified (Z-score normalised); shown in box- and-whiskers plot: 10–90 percentile; +, mean; dots, outliers; midline, median; boundaries, quartiles. n=2 independent experiments, 4 replicates/condition, minimum 500 cells/condition in total. p values: (Welsh’s t-test). **(F)** Intensity of Met antibody in membrane, cytoplasmic and juxtanuclear regions was quantified. Line graphs show relative region intensities compared to the intensity of the whole cell (Z-score normalised). n=2 independent experiments, 4 replicates/condition, minimum 500 cells/condition in total. p values: (Welsh’s t-test). **(G)** Schema, sub-cellular re-localisation of active Met upon HGF treatment. **(H)** PC3 cells expressing Scr, *IQSEC1 KD4, LRP1* or *SORL1* shRNA were stimulated with HGF. Cells were stained for phospho-Met (black), F-actin (green outlines) and Hoechst/nuclei (magenta). Magnified images of boxed regions are shown (lower panels). Arrows, reduction of pMet in peripheral regions. Cartoon, sub-cellular localization of active Met under different conditions. **(I-J)** Quantitation of spot intensity (Z-score normalised) is shown for images in **H** in box-and-whiskers plot: 10–90 percentile; +, mean; dots, outliers; midline, median; boundaries, quartiles. n=2 independent experiments, 4 replicates/condition, minimum 500 cells/condition in total. p-values (one-way ANOVA). **(K)** Schema, regulation of Met internalisation, but not recycling, by IQSEC1 and LRP1. All p values: n.s. not significant, *p≤0.05, **p≤0.01, ***p≤0.001 and ****p≤0.0001.

Paradoxically, IQSEC1-depleted cells displayed decreased total Met levels (Fig S6A,B), but an increased half-life of Met (Fig S6E), suggesting altered trafficking routes. We examined whether this was due to altered endocytosis and/or recycling of Met (Fig 5C). We developed an image-based analysis of Met trafficking using a fluorescently conjugated anti-Met antibody (Fig 5C, D). This allowed dual quantitation of internalisation levels and localisation to cellular sub-regions (membrane, cytoplasm, juxtanuclear). We detected some impact of IQSEC1 depletion on Met trafficking, though in all instances the magnitude of effect was modest. In control cells Met was efficiently labelled at the cell surface, appeared in peripheral endosomes after 10 min internalisation, then clustered in the juxtanuclear region by 30 min (Fig 5D-F). IQSEC1-depleted cells showed a modest but significant delay in the internalisation and transit of Met from the periphery to the juxtanuclear region (Fig 5D-F), representing a decrease in total Met internalisation levels. We observed no significant difference in recycling of internalised Met (Fig 5E, 10 min), but defects in internalisation were re-apparent in extended time points of recycling that allowed for re-internalisation (30 min). Thus, IQSEC1 is required for efficient internalisation, but not recycling, of Met (Fig 5F).

We examined whether the positive (LRP1) and negative (SORL1) interactors of IQSEC1-regulated invasion we identified may also function by controlling activated Met trafficking. LRP1/SORL1 are regulators of endocytic sorting of a number of transmembrane proteins ^50, 51^. In control cells, puncta containing activated Met (pY1234/5-Met, pMet) were distributed throughout the cytoplasm. Treatment with HGF induced a reduction in peripheral pMet puncta, concomitant with puncta clustering in the juxtanuclear region (Fig 5G-H), which was abolished by IQSEC1 depletion (Fig 5H-J). Consistent with a positive role for LRP1 in invasion, LRP1, but not SORL1, depletion also blocked redistribution of pMet away from the cell periphery in response to HGF (Fig 5H-J). These data reveal that IQSEC1 and LRP1 control Met-induced invasion by regulating Met transport to the juxtanuclear region (Fig 5K), an essential function for signalling from Met ^9^. The magnitude of effects of IQSEC1 depletion on trafficking, however, were modest. We therefore turned to potential effects on IQSEC1 in regulating signalling from Met.

### IQSEC1-ARF signalling controls phosphoinositide generation in invasive protrusion tips

We next examined whether the major effect of IQSEC1-ARF5/6 on Met signalling is to contribute to Akt activation. Generation of PI(3,4,5)P_3_ (PIP_3_) from PI(4,5)P_2_ is an essential event in the Akt signalling cascade, allowing PIP_3_-dependent recruitment of Akt to the cortex, PIP_3_-dependent PDPK1 phosphorylation of Akt T308 ^52^ and the PIP_3_-dependent release of SIN1 from RICTOR to allow Akt S473 phosphorylation ^53^. PIP_3_ generation requires the sequential action of PI4-kinases (PI4K), PIP5-kinases (PIP5K), and PI3-kinases (PI3K) to convert PI to PIP_3_. All ARF GTPases can recruit and activate PIP5Ks ^54–56^, the latter of which there are three isoforms, PIP5K1A-C (PIP5K1*α*/*β*/*γ*). ARF6 can activate PI3K signalling in melanoma ^57^ and function to recruit PIP5K1*α* to Met ^58^. We examined whether IQSEC1-ARF controls Akt activation by regulating PI(4,5)P_2_ production as a precursor to PIP_3_ generation (Fig 6A).

**Figure 6.**
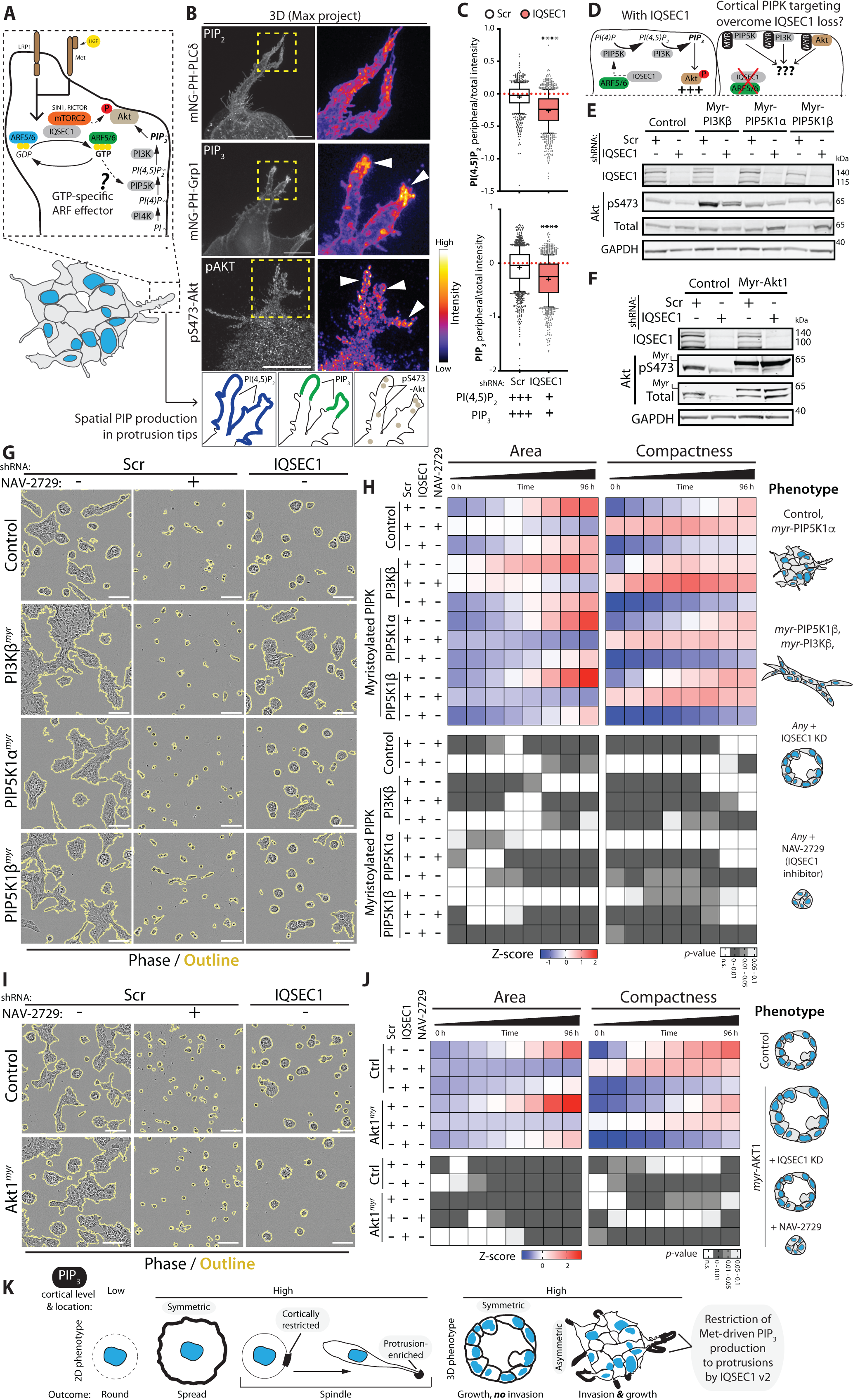
IQSEC1-ARF signalling controls phosphoinositide generation in invasive protrusion tips. **(A)** Schema, signalling pathways in protrusive tips of PC3 acini. **(B)** PC3 acini expressing mNeonGreen tagged PH-PLCδ or PH-Grp1 were fixed after 3 days. PC3 acini were also stained with phospho-S473 Akt antibody. FIRE LUT used to show localisation and intensity of mNeonGreen or pAkt. Magnified images of boxed regions are also shown. White arrowheads, localisation at protrusive tips. Scale bars, 10 µm. Cartoon, spatial PIP production in protrusive tips. **(C)** Quantitation of PI(4,5)P_2_ (upper panel) or PIP_3_ in the presence or absence of IQSEC1 is shown in box-and-whiskers plot: 10–90 percentile; +, mean; dots, outliers; midline, median; boundaries, quartiles. Values are peripheral/total intensity/cell. n=2 independent experiments, 4 replicates/condition, >500 cells/condition in total. p-values (one-way ANOVA): ****p≤0.0001. **(D)** Cartoon, PIPK targeting in presence or absence of IQSEC1. **(E)** Western blot of PC3 cells expressing Myr-FLAG-Cre (Control), Myr-FLAG-PIP5Kα, Myr-FLAG-PIP5Kβ or Myr-FLAG-PI3Kβ and either Scr or IQSEC1 shRNA (KD4). Anti-IQSEC1, phospho-S473 Akt, total Akt and GAPDH (loading control for IQSEC1 blot) antibodies were used. **(F)** Western blot of PC3 cells expressing Myr-FLAG-Cre (Control) or Myr-Akt1 and either Scr or *IQSEC1* KD4 shRNA using anti-IQSEC1, phospho-S473 Akt, total Akt and GAPDH (loading control for IQSEC1 blot) antibodies. **(G)** Phase images of PC3 acini described in **E** are shown. Scr acini were also treated with NAV-2729 (IQSEC1 inhibitor). Scale bars, 100μm. **(H)** Quantitation of images shown in **G**. Heatmaps show area and compactness measurements as Z-score-normalised values (upper panels). p-values (one-way ANOVA): greyscale values as indicated (lower heatmaps). n=3 independent experiments with 4 replicates per condition. 1,693 - 2,435 acini/condition quantified in total. Cartoon, acini phenotype representative of each condition. **(I)** Phase images of PC3 acini described in **F** after 96 hours. Scr acini were also treated with IQSEC1-inhibiting compound NAV-2729. Scale bars, 100μm. **(J)** Quantitation of images shown in **I**. n=2 independent experiments with 4 replicates/condition, 1,287-2,363 acini/condition in total. Cartoon, acini phenotype representative of each condition. **(K)** Schema, summarizes the relationship between location and level of peripheral PIP_3_ and 2D and 3D PC3 phenotype.

We investigated the relationship between cell shape and PIP level and localisation using biosensors and antibodies to each lipid. Cortical PI(4,5)P_2_ levels were not different between cell shape classes (spindle, spread, round) (Fig S7A, B). In contrast, cortical PIP_3_ was elevated in both spindle and spread shapes, but with alternate distributions. Spread cells had the highest PIP_3_ levels, which was uniformly cortical. In contrast, though spindle cells displayed lower cortical PIP_3_ than spread cells, PIP_3_ was instead enriched in the tips of protrusions in 2D and 3D (Fig S7A-C, 6B). This suggests that a combination of level and asymmetry of PIP_3_ promotes invasive function (Fig S7D).

To align the levels of PIPs with their distribution we validated that anti-PIP antibodies detected total cellular PIP levels, observing they are responsive to PI3K inhibition (LY294002) or activation (using myristoylated PI3K) (Fig S7E-G). In contrast to PI3K inhibition, which decreases total and cortical levels of PIPs (Fig S7F, H), IQSEC1 depletion lead to a striking alteration in PIP levels and distribution. IQSEC1-depleted cells displayed an overall increase in PI(4,5)P_2_ and PIP_3_ levels (Fig S7I), concomitant with a decrease in cortical PIPs (Fig 6C, S7C) and a redistribution of both PIPs from the cell cortex to intracellular compartments (Fig S7A). This reduction of cortical PIP_3,_ in the absence of IQSEC1 was observed equivalently whether the cell cortex was defined using either F-actin or a cytoplasmic stain (CellMask) (Fig S7J). Reduced cortical PIP3 levels in IQSEC1-depleted was comparable to the magnitude of PI3K-inhibited cells (Fig S7H). These data suggest that IQSEC1-ARF-PIP5K is not essential for general PI(4,5)P_2_-PIP_3_ generation, but rather that a LRP1-Met-IQSEC1 v2 complex spatially restricts PIP_3_ production to the tips of invasive protrusions (Fig 6A, S7D).

IQSEC1 v2 promotes single spindle cells (Fig S2D) and 3D invasion (Fig 2E-F), while IQSEC1 v4 promotes round shape. No difference in cortical PI(4,5)P_2_ distribution between cell shape classes (spindle, spread, round) was observed (Fig S7A, B). Accordingly, rescue of IQSEC1 depletion with either IQSEC1 v2 or v4 partially restored PI(4,5)P_2_ levels to those observed in control cells (Fig S7K). In contrast, the invasion-inducing IQSEC1 v2 supported more robust activation of PIP_3_ generation than could IQSEC1 v4 (Fig S7K). Given that IQSEC1 v2 induces a shape switch to spindle, and spindle cells have asymmetric cortical PIP_3_, this suggest that IQSEC1 v2-directed cortical asymmetry, rather than general production, of PIP_3_ drives invasion.

We next examined the components of PIP_3_ generation that drive invasion. Isotype-selective Class I PI3K inhibitors revealed that either PI3K*β* or PI3K*δ* inhibition decreased pAkt levels, while PI3K*α* or PI3K*γ* inhibition paradoxically increased pAkt levels (Fig S8A). However, only PI3K*β* or Akt inhibition significantly attenuated 3D growth and invasion (Fig S8B-D). Depletion of PIP5K1A-C revealed that PIP5K1*β* depletion showed the most robust decrease in growth, invasion, and pAkt levels (Fig S8E-G). These data suggest a PIP5K1*β*→PI3K*β*→Akt pathway for 3D growth and invasion.

Our data suggest that IQSEC1-ARF5/6 controls the PIP production and trafficking required for PIP5K1*β*→ PI3K*β*→ Akt signalling during invasion (Fig 6A). We tested whether generalised cortical targeting of this PIP-Akt pathway could overcome the need for IQSEC1 (Fig 6D). We targeted each of PIP5K1*β*, PI3K*β*, Akt1 to the cortex through addition of a myristoylation sequence, and inhibited IQSEC1 by shRNA or chemical means (NAV-2729) (Fig 6D-F).

Myristoylation of PI3K*β* (PI3K*β^myr^*) or PIP5K1*β* (PIP5K1*β^myr^*), but not PIP5K1*α* (PIP5K1*α^myr^*), increased Akt phosphorylation, 3D growth and invasion, with the most robust increase occurring upon PI3K*β^myr^* expression (Fig 6E-H). As expected in control cells, IQSEC1 depletion reduced pAkt levels and attenuated 3D growth and invasion (Fig 6E-H). Myristoylation of PI3K*β* (PI3K*β^myr^*) or PIP5K1*β* (PIP5K1*β^myr^*), but not PIP5K1*α* (PIP5K1*α^myr^*), increased 3D growth and invasion with the most robust increase occurring upon PI3K*β^myr^* expression (Fig 6E, G, H). Strikingly, IQSEC1 perturbation (shRNA, NAV-2729) had a disproportionate effect on invasion versus growth. While growth was blunted, in IQSEC1-perturbed conditions invasion was abolished (Fig 6E, G, H). NAV-2729 treatment blocked the ability of all cells in 3D to become acini. Single cells initially increased slightly in area and adopted irregular shapes, manifested in increased compactness, but not in invasion. Thus, generalized cortical recruitment of PIPK signalling is sufficient to drive growth, but not invasion. Invasion requires IQSEC1-dependent enrichment of signalling to invasion tips (Fig 6K).

### IQSEC1-ARF regulates localization and activation of Akt

Akt signalling showed a different influence on 3D behaviours than that observed for PI3K*β* or PIP5K1*β*. Our data suggest that IQSEC1 may contribute to Akt S473 levels by at least two mechanisms: influencing PIP5K1*β*-directed PI(4,5)P_2_ production, and interaction with RICTOR-SIN1 (Fig 6A). While IQSEC1 depletion lowered endogenous pAkt levels, expression of myristoylated Akt1 (Akt1*^myr^*) overcame the requirement of IQSEC1 for pS473 phosphorylation (Fig 6F, I, J). As predicted, general cortical targeting of Akt1*^myr^* robustly increased 3D area compared to control cells, but not invasion (Fig 6I, J). Strikingly, although Akt1*^myr^* no longer required IQSEC1 for phosphorylation events normally indicating ‘activeness’ (i.e. pS473-Akt, Fig 6F), IQSEC1 depletion still reversed the Akt1*^myr^*-induced 3D growth increase to levels below control cells, and abolished invasion; NAV-2729 was similarly an inhibitor of these processes (Fig 6I, J). Thus an effector of PIP_3_ signalling, Akt, may also require IQSEC1-dependent asymmetry at the cortex for full signalling output.

As experimentally ‘active’ Akt (asymmetrically cortically targeted Akt1*^myr^*; as defined by pS473 levels) was unable to induce oncogenic signalling in IQSEC1-depleted cells we examined whether this was due to altered Akt localisation. In 2D, pAkt localised to cortical regions and to puncta distributed through the cytoplasm (Fig S8H). In 3D, a pool of pAkt was enriched in protrusion tips (Fig 6B, white arrowheads), as well as to puncta throughout the protrusion and acinus body. In contrast, IQSEC1-depleted cells displayed an increased cortical pAkt aggregate size, but with strongly reduced intensity (Fig S8H-K). These data reveal that IQSEC1 is required for the signalling from, and asymmetry of, the PIP_3_ effector Akt.

### IQSEC1 regulates growth and invasion *in vitro* and *in vivo*

Our data indicate that LRP1-Met-IQSEC1-promoted enrichment of ARF5/6-dependent PIP_3_ signalling to the tips of cellular protrusions induces invasion rather than growth. We tested the generalizability of IQSEC1 inhibition to inhibit growth and invasion mechanisms in commonly used prostate cancer models. mRNA expression and western blotting indicated that, with the exception of LRP1 in 22Rv1, all components of the LRP1-Met-IQSEC1-ARF5/6 pathway are expressed in examined prostate cancer cell lines (Fig S9A, B).

Chemical inhibition (NAV-2729, Fig S9C-H) or genetic depletion (Fig S10A-F) of IQSEC1 attenuated growth and/or invasion in a range of 3D cancer cell models. This included upon ectopic overexpression of the LRP1-IQSEC complex in cells with low endogenous levels (LRP1^TM^-GFP in 22Rv1, GFP-IQSEC1 v2 in DU145), HGF treatment of the mixed morphology DU145 cultures to resemble the spindle-type invasion of PC3 (Fig S9, S10), highly invasive human breast cancer cells (MDA-MB-231), murine pancreatic ductal adenocarcinoma cells (PDAC; KC-PTEN, K-rasG12D/PTEN-null^59^) and patient derived PDAC cells (TKCC-07) (Fig S10A, C-F). Thus, IQSEC1 is required for growth and invasion across a number of 3D cell models from different cancer types.

We examined the *in vivo* role of IQSEC1 by intraprostatic xenograft of IQSEC1-depleted PC3 cells (Fig 7A). While IQSEC1 depletion did not significantly attenuate tumour incidence, tumour area and volume were significantly reduced (Fig 7B-E). As predicted from our 3D *in vitro* studies, metastatic activity was strongly decreased in IQSEC1-depleted cells. Both the incidence and number of macrometastases were significantly decreased in IQSEC1-depleted cells (Fig 7F, G). Wide-spread dissemination of macrometastases was observed in controls cells. In the few mice presenting macrometastases in IQSEC1-depleted cells, these were limited to prostate proximal lymph nodes (with the exception of a singular diaphragm-located tumour) (Fig 7H), and showed no difference in proliferation or apoptosis to controls, confirming a bona fide effect on movement (Fig 7I, J). These data indicate an essential requirement for IQSEC1 in metastasis *in vivo*.

**Figure 7.**
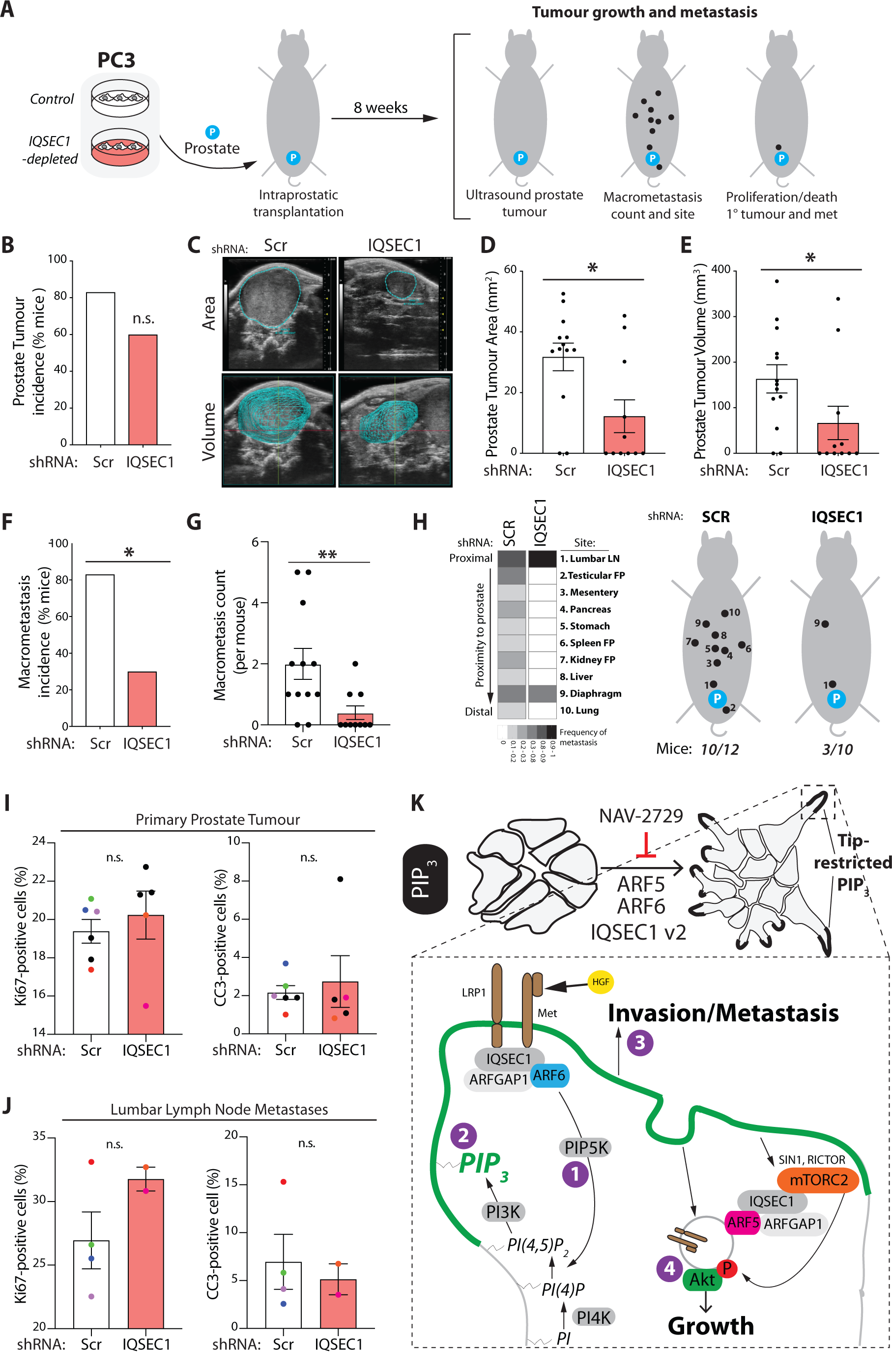
IQSEC1 regulates growth and invasion *in vivo* and IQSEC1-ARF signalling is associated with poor patient outcome. **(A)** Schema, intraprostatic transplantation of PC3 cells expressing either Scr (control, 12 mice) or *IQSEC1* KD4 shRNA (10 mice) into CD1-nude male mice. Tumour growth was determined after 8 weeks using ultrasound and the incidence and location of macromets counted. **(B)** Bar graph shows prostate tumour incidence (% mice). p-values (Chi-squared): n.s. not significant. **(C)** Ultrasound images show area and volume of representative prostate tumours in mice injected with PC3 cells expressing either Scr or IQSEC1 KD4. **(D-E)** Area **(D)** and volume **(E)** of each prostate tumour detectable by ultrasound was measured. p-values (Mann-Whitney): *p≤0.05. Note that one mouse in each condition (Scramble, IQSEC1 KD) had two primary prostate tumours. Control, 12 mice, 13 tumours. *IQSEC1* KD4 shRNA (10 mice, 11 tumours). **(F-G)** Macrometastasis incidence was counted and is presented as % mice. **(F)** and count per mouse **(G).** p-values (Chi-squared and Mann-Whitney respectively): *p≤0.05, **p≤0.01. **(H)** Heatmap shows the frequency at which metastasis occurred at locations with different proximity to the prostate. Cartoon, summarizes the locations metastasis was observed. **(I-J)** Tumour **(I)** and lymph node **(LN) (J)** sections were stained with anti Ki67 and Cleaved Caspase 3 (CC3) antibodies and the percentage of positive cells quantified. Each mouse is represented by a differently coloured point on the bar graphs. p-values: (Mann-Whitney) n.s. not significant. **(K)** Schematic depicting the signalling mechanisms involved in IQSEC1 dependent model of prostate cancer growth and invasion.

Elevation in IQSEC1, ARF5 and ARF6 levels was associated with clinical outcome in prostate cancer across 12 studies, representing 2,910 patients (Fig 8A). Increased Copy Number (CN) of IQSEC1 occurred most frequently in advanced prostate cancers (Fig 8B, CR, castrate-resistant; NE, neuroendocrine), associated with a gain in 3p Status (Fig 8C; IQSEC1, 3p25.2-25.1). Primary prostate tumours with elevated IQSEC1 were of significantly higher grade (Fig 8D), were associated with tumour-bearing lymph node positivity (Fig 8E) and metastases (Fig 8F). Mirroring the ablation of wide-spread metastasis of IQSEC1-depleted PC3 xenografts (Fig 7H), IQSEC1 increase occurred most frequently in samples from diverse metastatic sites, but particularly bone, liver and lymph node (Fig 8G). IQSEC1 increase occurred exclusively with Androgen Deprivation Therapy, retained presence of tumour after therapy, and increased levels of the Androgen Receptor V7 variant, a major mechanism for escape from androgen deprivation (Fig 8H-J). A clinical indicator of disease, serum PSA levels, was elevated in patients displaying combined IQSEC1 and ARF5/6 elevation (Fig 8K). This suggests a clinical association of IQSEC1-ARF5/6 gain with therapy resistance and metastasis.

**Figure 8.**
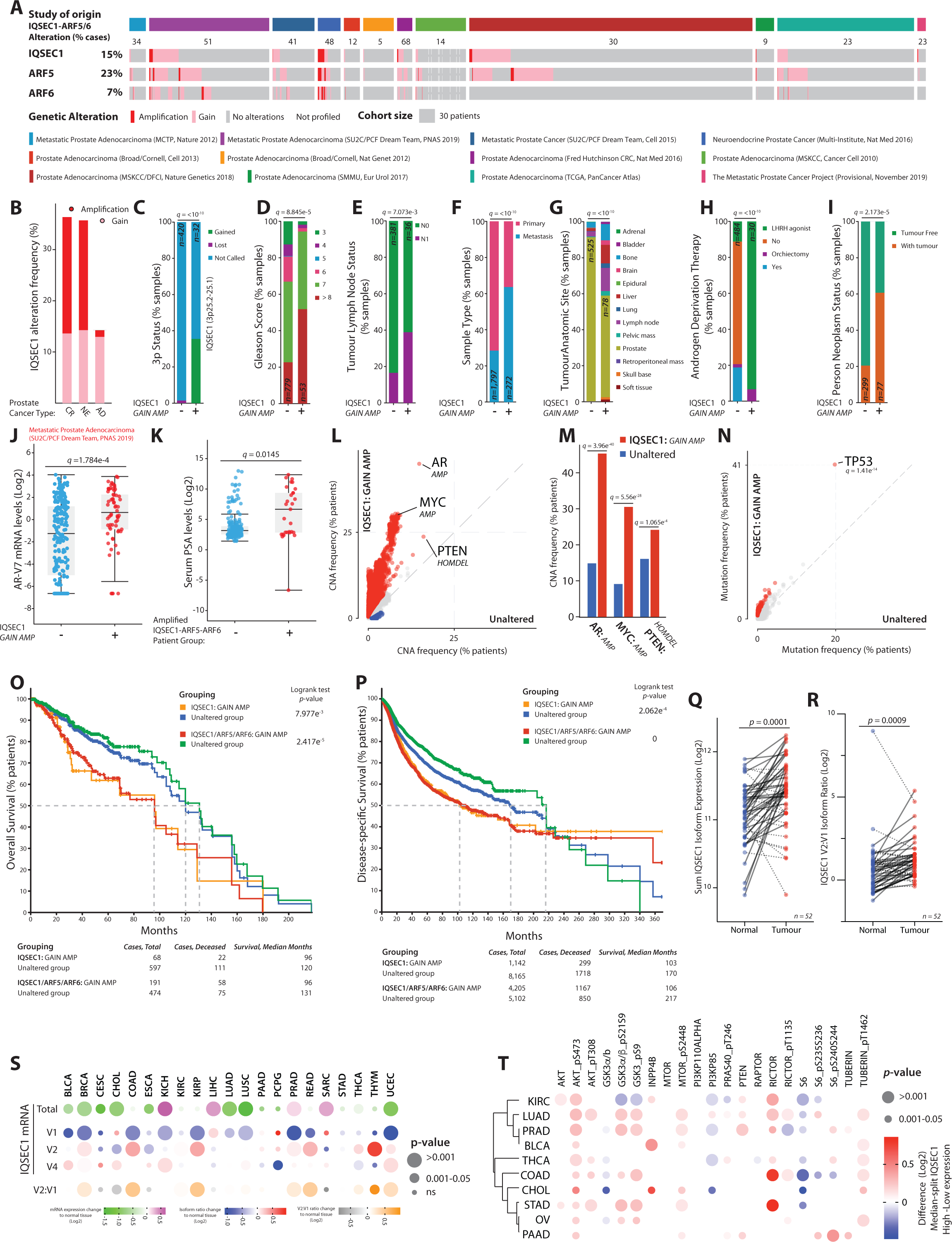
IQSEC1 is associated with metastasis, treatment resistance, and poor clinical outcome. **(A)** CN increase frequencies (percentage of patients/cohort) in IQSEC1, ARF5, ARF6 across prostate cancer cohorts for subsequent analyses. **(B)** CN increase frequencies across prostate cancer types from cohorts in A. AD, prostate adenocarcinoma; NE, neuroendocrine; CR, Castrate-resistant. **(C-J)** Comparison of clinical metrics, represented as percentage of samples, between IQSEC1 CN-amplified patients (+) compared to non-amplified comparison group (-) for **(C)** 3p Status (-, n=420; +, n=32) (note IQSEC1 proband, 3p25.2-25.1), **(D)** Gleason Score (-, n=779; +, n=53), **(E)** Tumour Lymph Node Status. N0, no lymph node positivity. N1, lymph node positivity. (-, n=381; +, n=36), **(F)** Sample Type (-, n=1,797; +, n=272), **(G)** Tumour Anatomic Site (-, n=525; +, n=78), **(H)** Androgen Deprivation Therapy. Luteinizing hormone-releasing hormone, LHRH. (-, n=484; +, n=30), **(I)** Person Neoplasm Status (-, n=299; + n=77), and **(J)** Androgen Receptor (AR) isoform V7 mRNA levels (Log2) (-, n=221; + n=69). P values, (C-I) Chi-squared Test or (J) Kruskal Wallis Test; Q values, Benjamini-Hochberg adjustment. **(K)** Serum levels of Prostate-specific antigen (PSA) (represented as Log2) in patients with IQSEC1, ARF5 and IQSEC1 CN increase. Patients, Altered, n=22; Unaltered, n=86. Q value, Kruskal Wallis Test. **(L-N)** Differential Copy Number Alteration **(L, M)** or **(N)** mutation frequencies between IQSEC1 CN-amplified patients compared to non-amplified comparison group. Red, IQSEC1 CN increase-associated. Blue, control group-associated. Total patients n=3,157, unaltered, n=2308, altered=849. Q-values, one-sided Fisher exact test with Benjamini-Hochberg adjustment. **(O-P)** Overall patient survival (percentage of patients across months) across two groupings compared to unaltered patients: i) IQSEC1 CN increase or ii) combined IQSEC1, ARF5, ARF6 CN increase for **(O)** prostate cancer patient cohorts in A (n=665) or **(P)** across the pan-cancer TCGA cohorts (n=9,307). P-values, Log-rank T-test. **(Q-R)** Matched normal tissue (Normal) and primary prostate tumour (Tumour) IQSEC1 mRNA levels for **(Q)** sum of IQSEC1 isoforms or **(R)** ratio of IQSEC1 v2 to V1. Log2-normalised values. Blue, normal tissue. Red, tumour tissue. Unbroken line, increased value in tumour; dotted line, decreased value in tumour, n=52. P-values, Independent Groups T-test. **(S)** Bubble heatmap of IQSEC1 expression in tumour compared to normal tissue across pan-cancer TCGA cohorts. Values are Log2 normalized fold change compared to normal tissue. Colouring, total IQSEC1 expression (green to magenta), individual IQSEC1 isoforms (V1, V2, V4; blue to red), or V2-to-V2 ratio (grey to orange). Circle size, p-value. Patient sample size (n): Normal Tissue, N, 643; T, Tumour Tissue, T, 6,716. See Table S2 for breakdown by tumour type. P-values, Independent Groups T-test. **(T)** Bubble heatmap of protein expression in the PI3K-AKT signalling pathway from RPPA analysis of pan-cancer TCGA cohorts divided into High versus Low IQSEC1 expression. Values are Log2-transformed difference between a median split of total IQSEC1 mRNA levels. Red, co-occurring with high IQSEC1; blue, associated with low IQSEC1 levels. Only cancer types presenting increased AKT_pS473 are included. Row sorting based on dendrogram. Circle size, p-value. P-values, Independent Groups T-test.

We examined the association of IQSEC1, ARF5, and ARF6 elevation with frequent genomic alterations in prostate cancer. IQSEC1 increase was associated with amplifications particularly in AR and MYC and to a lesser extent with loss of PTEN, and prominently with TP53 mutation, but not common gene fusion events (Fig 8L-N, S11A). ARF6 increase followed a similar profile, while ARF5 increase was associated with a broader range of mutational events. IQSEC1 allowed robust stratification of prostate cancer patients, stratifying a 24-month decrease in median overall survival in IQSEC1-elevated patients (Fig 8O, 665 patients). Inclusion of ARF5/6 increase did not change the median overall survival of this group, but rather extended the median survival of the control arm, resulting in a 35-month survival difference compared to the IQSEC1-ARF5-ARF6 elevated group. IQSEC1 increase was similarly associated with TP53 mutation and MYC amplification across the pan-cancer The Cancer Genome Atlas (TCGA) dataset, representing 10,449 patients (Fig S11A-D). Strikingly, IQSEC1, ARF5 and ARF6 increase similarly stratified patient survival when all tumour types were considered together, providing an exceptional >9-year (111 months) median survival increase for non-IQSEC1-ARF5/6-amplified patients (across 9,307 patients) (Fig 8P).

Comparison of matched normal-tumour tissue from the TCGA prostate cohort (n=52) showed a significant increase in overall IQSEC1 mRNA in tumours (Fig 8Q). This was accompanied by a switch from the non-invasive IQSEC1 v1 isoform, to the pro-invasive IQSEC1 v2 in tumours (Fig 8R). Analysis of the pan-cancer TCGA dataset for tumour types with profiled normal tissue revealed that four tumour types (Kidney, KICH; Liver, LIHC; Prostate; Sarcoma, SARC) displayed IQSEC1 mRNA higher than normal tissue (Fig 8S). However, seven tumour types showed that rather than an increase in overall IQSEC1, instead tumours underwent an isoform switch from V1-to-V2 (Fig 8S). This demonstrates that isoform switching to IQSEC1 v2 is a major event associated with tumourigenesis in patients.

Finally, we examined the association of IQSEC1 mRNA levels across the pan-cancer TCGA dataset with common tumour-associated signalling pathways from Reverse Phase Protein Array (RPPA) data. Within each cancer type IQSEC1 expression was divided into high and low using a median split of mRNA levels, and protein differences between each group that consistently and significantly trended in the same direction across a quarter of all cancer types was calculated (Fig S11E). Mirroring IQSEC1 regulation of phosphoinositide signalling in PC3 cells, a clear PI3K-AKT signature was associated with high IQSEC1 levels across ten tumour types (Fig 8T). Together, these data indicate a role for tumour-associated isoform switching to IQSEC1 v2 to promote PI3K-AKT signalling and metastasis across cancer types, resulting in treatment resistance and a robust decrease in patient survival.

## Discussion

Our results clarify a long-held conundrum of how the same signalling pathways lead to distinct biological outputs: while a uniform peripheral signal induces growth, spatially enriching phosphoinositide metabolism downstream of an RTK causes local remodelling of peripheral subdomains into invasive protrusions.

We describe that IQSEC1 is a key node in the spatial enrichment, rather than total amount, of PIP_3_ signalling to promote invasion. Our data support a model whereby pro-invasive IQSEC1 isoforms form a complex with the HGF receptor Met and the endocytic receptor LRP1 at membranes that will develop into invasive protrusions (Fig 7K). IQSEC1 activates GTP loading on ARF6 to 1) stimulate PIP5K-mediated peripheral PI(4,5)P_2_ production, which is 2) a precursor to PI3K-mediated PIP_3_ production to 3) promote invasive protrusion formation. Production of PIP_3_ at invasive protrusions also triggers cell growth, concomitant with internalisation of Met and 4) activation of pAkt downstream of the mTORC2 complex. Met and pAkt are retrograde transported in the body of protrusions on ARF5 endosomes to a juxtanuclear signalling compartment, essential for growth and invasion. Inhibiting this IQSEC1-mediated ARF5/6 activation abolishes the collective growth and invasion/metastasis *in vitro* and *in vivo*.

Multiple isoforms of IQSEC1 exist through combination of alternate start sites and alternate splicing, though the regulators of these events are unknown. Such N- and C-terminal extensions operate in a hierarchy; N-terminal extensions provide association with endocytic co-receptors LRP1 and SORL1, while alternate C-termini can either positively or negatively enhance invasive activity. SORL1 and LRP1 are members of the LDLR family, co-receptors involved in endocytic control of a milieu of processes ^42^. SORL1 and LRP1 shared the same N-termini binding region, were anti-correlated in their expression between invasive and non-invasive cells, and possessed antagonistic effects on invasion. This supports a notion that the function of IQSEC1 is highly contextual, dependent on not only which IQSEC1 transcript is expressed, but also the repertoire of potentially competitive interactors expressed. It is notable switching to the pro-invasive IQSEC1 v2 occurs in at least a quarter of all sampled tumour types, suggesting this may be a major mechanism underpinning tumourigenesis.

LRP1 is a multifunctional endocytic receptor involved in a number of biological functions ^42^. We report this as a crucial pro-invasive partner of IQSEC1 and Met. Endogenous LRP1 and Met were found in a complex irrespective of IQSEC1, suggesting that IQSEC1 binds an existing complex to couple their endocytic itinerary. This likely also requires the function of the extracellular cleaved 515kDa fragment of LRP1, as the transmembrane fragment alone is insufficient to drive invasion when expressed ectopically. LRP1 may help generate PIP_3_ during invasion through its ability to form an LRP1-Rab8a:GTP-PI3K complex, to control Akt signalling on endosomes, as it does in other contexts ^60^.

Protrusive invasion requires a discrete pool of protrusive activity concomitant with global inactivity in surrounding regions, akin to pushing out from single point inside a balloon rather than simultaneously from all surfaces. IQSEC1 v2 localised with Met and LRP1 and coincided with PIP_3_ production at such a discrete event: the tips of invasive protrusions. Such exquisite spatial enrichment coupled to invasive protrusion may explain why the most prominent peripherally recruited IQSEC1 isoform (v4) fails to induce robust invasion, and perhaps why we see significant, but modest global GTP loading changes in ARF5/6 upon IQSEC1 depletion. The identity of invasion-specific PIP_3_ effectors remains to be demonstrated, but one likely candidate is Rac1, which is dependent on PIP_3_ for activation ^61^.

Why are both ARF5 and ARF6 required? Although ARF5 phenotypes most closely mimicked that of IQSEC1, there was still a dependency on ARF6. ARFs can act in pairs ^61^, and our data suggests that ARF5/6 may act at sequential steps: ARF6 at the cortex, and ARF5 in endosomes. It is interesting to note that previously described fast-cycling mutants of either ARF ^36^ were unable to induce collective morphogenesis like WT ARFs, suggesting that precise control over GTP cycling is essential for collective invasion. Indeed, a key effector identified was ARFGAP1, which in our hands functioned as an effector rather than an antagonist of ARF signalling, similar to the dual effector terminator function described for other ARFGAPs ^47–49^. That this GAP associates with the GEF supports the notion of tight control over GTP cycling. Whether ARF5/6 both, or separately, interact with the two identified effectors (ARFGAP1, PIP5K1*β*), is unclear.

The ARF GTPase-dependent recycling of Met promotes sustained ERK signalling required for migration in other cells ^58, 62^. Internalisation of Met is key to full oncogenic output ^8, 9^. IQSEC1 has modest effects on internalisation and juxtanuclear trafficking of Met, suggesting the major role of IQSEC1 is to couple Met to its downstream component Akt. Met recycling was not perturbed in IQSEC1-depleted cells, suggesting another ARFGEF may promote this function. The PI(4,5)P_2_/PIP_3_-dependent Cytohesin-1 ARFGEF is one candidate for this, as we note upregulation in PC3, and it is required downstream of Met for invasive activity in other cells ^63^. Cytohesin-1 alternate splicing controls its recruitment to membranes by controlling differential affinity for PI(4,5)P_2_ or PIP_3_ ^64, 65^. Such a GEF would likely act subsequently to IQSEC1-directed control of peripheral PI(4,5)P_2_ levels. Thus, IQSEC1 may sit at the top of a cascade of ARF GEFs controlling Met endocytosis, recycling, and signal output. This may explain why IQSEC1 inhibition alone can reverse ARF5/6 overexpression, despite co-expression of numerous ARFGEFs in PC3 cells.

A striking result is that although IQSEC1-ARF5/6 would classically be considered ‘upstream’ of PI(4,5)P_2_, PIP_3_, and Akt, inhibition of IQSEC1 nonetheless counteracts the experimental activation of this pathway at multiple levels. Our data suggests that this is due to IQSEC1 controlling the spatial enrichment of these signals. This suggests targeting mechanisms of spatial enrichment of signalling pathways may be effective in inhibiting oncogenic signalling, as previously demonstrated for c-Met ^8^, and demonstrated here for IQSEC1 and the PIP_3_-Akt pathway.

Critically, we demonstrate that IQSEC1 inhibition blocks growth and invasion in a multitude of *in vitro* 3D models across several cancer types, and tumour growth and metastasis *in vivo*. Elevated IQSEC1 expression is associated with clinical metrics of poor outcome: higher-grade tumours, metastasis, treatment resistance, PI3K pathway activation. Consequently, IQSEC1-ARF5/6 stratifies long-term recurrence-free survival in prostate cancer and more generally across a number of tumour types. In conclusion, we identify a molecular mechanism that allows cells to determine whether to grow or invade in response to activation of an RTK: enrich signalling output, such as PIP_3_ production, to peripheral subdomains to convert these into invasive protrusion, over a generalised peripheral PIP_3_ production which will cause a growth response.

## Methods

### Cell culture

PC3 (ATCC), PC3 E-Cad+, TEM4-18, TEM2-5, GS689.Li, GS694.LAd, GS683.LALN, JD1203.Lu, GS672.Ug (M. Henry, University of Iowa), PC3-Epi and PC3-EMT (K. Pienta, Johns Hopkins School of Medicine) and LNCaP cells (H. Leung, Beatson Institute) were cultured in RPMI-1640 supplemented with 10% Fetal Bovine Serum (FBS) and 6mM L-glutamine. RWPE-1, RWPE-2, WPE-NB14 and CA-HPV-10 cell lines (ATCC) were grown in Keratinocyte Serum Free Media (K-SFM) supplemented with 50μg/ml Bovine Pituitary Extract (BPE) and 5ng/ml Epidermal Growth Factor (EGF). DU145 cells (ATCC) were maintained in Minimum Essential Medium (MEM) supplemented with 10% FBS and 6mM L-glutamine. VCaP, MDA-MB-231 (ATCC), murine KC Pten^fl/+59^ (J. Morton, CRUK Beatson Institute) and TKCC-07 cells were cultured in Dulbecco’s Modified Eagle Medium (DMEM) supplemented with 10% FBS and 6mM L-glutamine. The TKCC-07 cells used in this project were provided by the Australian Pancreatic Cancer Genome Initiative (APGI) at the Garvan Institute of Medical Research (www.pancreaticcancer.net.au). 22Rv1 cells (H. Leung, Beatson Institute) were grown in phenol free RPMI-1640 containing 10% charcoal stripped FBS and 6mM L-glutamine. HEK293-FT (Thermo Fisher Scientific) were cultured in DMEM with 10% FBS, 6mM L-glutamine and 0.1mM Non-Essential Amino Acids (NEAA) (all reagents from Thermo Fisher Scientific).

Growth factors or inhibitors were added as follows; 50ng/ml Hepatocyte Growth Factor (HGF) (PeproTech), 10μM NAV-2729 (Glixx Laboratories), 10μM QS11 (Tocris), 100μM chloroquine (CST), 20μM SecinH3 (Tocris), 25μM cycloheximide (Sigma), 1μM LY-294002 (Merck), 1μM AZD8835 (AstraZeneca), 0.1μM AZD8186 (AstraZeneca), 1μM AS-6052-40 (Stratech), 1μM Cal-101 (Stratech), and 10μM AktII (Calbiochem) inhibitors.

Cells were routinely checked for mycoplasma contamination. PC3, RWPE-1 and RWPE-2 cells were authenticated using short tandem repeat (STR) profiling.

### Generation of stable cell lines

Cell lines were made by co-transfecting plasmids with lentiviral packaging vectors (VSVG and SPAX2) into packaging cells (HEK293-FT) using Lipofectamine 2000 (Thermo Fisher Scientific). Viral supernatants were collected; filtered using PES 0.45μm syringe filters (Starlab) to remove cell debris, and concentrated using Lenti-X Concentrator (Clontech) as per the manufacturer’s instructions. Cells were then transduced with the lentivirus for 3 days before either FACS sorting or selection with 2.5μg/ml puromycin, 300μg/ml G418 (both Thermo Fisher Scientific), 10μg/ml blasticidin (InvivoGen) or 200μg/ml hygromycin (Merck). Stable knockdown of proteins was achieved using pLKO.1-puromycin, pLKO.1-hygromycin or pLKO.1-membrane tagged Venus (substituted for puromycin) lentiviral shRNA vectors. Sequences as follows: shScr (5’CCGCAGGTATGCACGCGT3’), IQSEC1 KD1 (5’GATCTATGAACGGATCCGTAA3’), IQSEC1 KD4 (5’CCAGTACCAGATGAACAAGAA3’), IQSEC1 KD murine (5’CCAGTGTTACTGTTGGCAAATCTC3’), Arf 5_1 (5’GATGCAGTGCTGCTGGTATTT3’), Arf 6_3 (5’GCTCACATGGTTAACCTCTAA3’), LRP1_1 (5’GATGCCTATCTGGACTATATT3’), LRP1_2 (5’GATCCGTGTGAACCGCTTTAA3’), MET_2 (5’GTGTGTTGTATGGTCAATAAC3’), SORL1_1 (5’GCCCAGTTTGTCACAAGACAT3’), SORL1_2 (5’CCTATGCCATTGCTGTCTTTA3’), RICTOR_1 (5’ACTTGTGAAGAATCGTATCTTCTC3’), RICTOR_2 (5’GCAGCCTTGAACTGTTTAA3’), SIN1_2 (5’CTAAGCAATCACGACTATAAA3’), ARFGAP1_1 (5’GTGCAGGATGAGAACAACGTT3’), ARFGAP1_2 (5’GCCAGTTCACTACGC AGTATC3’), PIP5K1α (5’GCTTCCAGGATACTACATGAA3’), PIP5K1β (5’ACGACAGGCCTAC ACTCTATT3’) and PIP5K1γ (5’CAACACGGTCTTTCGGAAGAA3’).

GFP-IQSEC1 v2 and GST-GGA3-GAT were kind gifts from J. Casanova (University of Virginia). All RNAi resistant variants and chimeras were made by mutagenesis or sub-cloning using fragment synthesis (GeneArt). GFP ARFs were kinds gift from P. Melancon (University of Alberta) and alternate fluorescent tags and mutations generated by sub-cloning. EGFP-PH-GRP1 and EGFP-PH-PLCδ were described previously^66^ and sub-cloned into mNeonGreen. Myr-FLAG-Cre, Myr-FLAG-PIP5Ks, Myr-FLAG-PI3Ks, Myr-Akt1 and RFP-GGA1-NGAT were purchased from Addgene. LRP1-EGFP was a kind gift from S. Kins (Technical University Kaiserslautern). Plasmids will be deposited at Addgene.

### IQSEC1 variant information

The nomenclature for IQSEC1 variants has been complicated by alternate names for IQSEC1 in literature and changing isoform designations at NCBI. Unification of nomenclature is presented in Table S3.

### Live 3D culture and analysis

Culture of cell lines as 3D acini was adapted from previous protocols ^5^. Briefly, single cell suspensions were made (1.5 x 10^4^ cells per ml) in the appropriate medium supplemented with 2% Growth Factor Reduced Matrigel (GFRM; BD Biosciences). 150μl of this mix was plated per well in a 96 well ImageLock plate (Essen Biosciences) pre-coated with 10μl of GFRM for 15 minutes at 37°C. Plates were incubated at 37°C for 4 hours, then imaged using an IncuCyte® ZOOM (Essen BioScience). Images were taken every hour for 4 days at 2 positions per well using a 10x objective lens. Sample size (n) and replicate number are stated for each experiment in Figure Legends. Outlines of phase and GFP-positive (where appropriate) acini were generated using a custom pipeline in CellProfiler. A custom macro in Fiji software was then used to colour code images from each time point, progressively coloured along a blue-to-red rainbow timescale, and concatenate them into one image per 12-hour block to reduce data dimensionality of multiday imaging. Using CellProfiler, measurements such as Area and Compactness, which could reliably measure size and protrusiveness of 3D PC3 objects respectively, were generated for each 12-hour block. Custom pipelines designed in KNIME Data Analytics Platform were then used to collate data from multiple experiments, normalise to controls, calculate Z-score and perform statistical analysis using one-way ANOVA. Normalised data and p-values are presented as heatmaps generated in PRISM 7 (GraphPad). Total number of acini quantified per condition is stated in each Figure Legend.

### 2D and 3D acini immunofluorescence and imaging

3D acini were set up as described above in either 8-well coverglass chamber slides (Nunc, LabTek-II) or in 96 well plates (Greiner) pre-coated with 60μl or 10ul GFRM respectively for 3 days. For 2D, cells were plated on a 96 well plate (Greiner) for 48 hours prior to fixation. Samples were washed twice with PBS and fixed in 4% paraformaldehyde for 15 minutes. Samples were then blocked in PFS (0.7% fish skin gelatin/0.025% saponin/PBS) for 1 h at RT with gentle shaking and primary antibodies added overnight (1:100 unless stated otherwise) at 4°C. After three washes in PFS, Alexa Fluor secondary antibodies (1:200), HCS CellMask™ Deep Red Stain (1:50000) or Hoechst (1:1000) (all Thermo Fisher Scientific) were added for 45 minutes at room temperature (RT). Samples were maintained in PBS, after 3 five minute washes in PBS, until imaging was carried out.

Cells or acini on chamber slides were imaged using a Zeiss 880 Laser Scanning Microscope with Airyscan. Images taken in super resolution mode on the Airyscan microscope were processed using the Zeiss proprietary ZEN software, exported as TIFF files and processed in Fiji. Cells or acini on 96 well plates were imaged using an Opera Phenix™ High Content analysis system and where appropriate Harmony High-Content Imaging and Analysis Software (PerkinElmer) was used to perform machine learning.

Antibodies used; Alexa Fluor 488, 568 or 647 phalloidin (Thermo Fisher Scientific), anti-IQSEC1 (Sigma, G4798), anti-IQSEC1 (Caltag-Medsystems, PSI-8009), anti-LRP1 (Sigma, L2295), anti-Met (CST, 3127), anti-Met phospho 1234/1235 (CST, 3077), anti-Akt phospho S473 (CST, 3787), DYKDDDDK Tag (CST, 14793, 1:800), anti-Ki67 (Thermo Fisher Scientific, 18-0192Z), anti Cleaved Caspase 3 (CST, 9661), anti-PtdIns(3,4,5)P3 and anti-PtdIns(4,5)P2 (Echelon Biosciences, Z-P345B and Z-P045).

### 2D and 3D Phenotypic analysis of fixed samples

Harmony High-Content Imaging and Analysis Software (PerkinElmer) was utilized to perform machine learning using a number of custom designed pipelines. For 2D morphology assays the shape of each cell was defined by either F-actin or HCS CellMask staining and machine learning then used to classify cells as either spindle, spread or round phenotypes. A custom pipeline was generated using KNIME Data Analytics Platform to collate data, calculate the log2 fold change of each phenotype over control and to calculate statistical significance using one-way ANOVA. In some cases this analysis was carried out in sub-populations of cells that either expressed a fluorescently tagged plasmid or were stained with a specific antibody.

The use of the nuclear, cytoskeletal and cytoplasmic reagents described above also allowed each cell to be segmented into specific sub-cellular regions i.e. nuclear, cytoplasmic and peripheral (defined as 5 pixels from outer edge of cell). When used in combination with fluorescently tagged proteins (such as reporters for phosphoinositides) or antibodies the mean intensity of a specific protein per cell (intensity/cell) or per sub-cellular region per cell (i.e. mean peripheral intensity/mean total intensity/cell) could be measured. Where appropriate ‘spots’ of positive staining were detected within these sub-cellular regions to more accurately calculate the mean intensity/cell or mean spot area/cell.

Custom KNIME pipelines described previously ^66^ were then used to collate data, to calculate the ratio over control and to determine statistical significance. In addition, these pipelines were adapted to measure total intensity of specific antibodies in 3D acini. Data are presented in box and whiskers plots as z-score normalized or as log2 fold change over control where each point represent one cell. Sample number (n), number of cells/acini imaged per experiment, statistical test performed and significance is stated in the appropriate Figure Legend.

Harmony High-Content Imaging and Analysis Software was also used to quantify the colocalization between mNeonGreen-tagged ARF5 or ARF6 and RFP-GGA1-NGAT. PC3 cells stably expressing ARF/GGA1 were FACS sorted and a population that was positive for both mNeonGreen and RFP selected. These cells were plated for 24 hours then treated with either NAV-2729 or QS11 overnight. Cells were then fixed, stained with CellMask and Hoechst and imaged using an Opera Phenix™. A custom pipeline was created to identify red (GGA1 +) and green (ARF +) ‘spots’ throughout each cell and to calculate the percentage of green spot area that overlapped with red spot area per cell. n=2 with 4 replicates per condition with > 500 cells imaged per condition.

### Animal studies

Animal experiments were performed under the relevant UK Home Office Project Licence (70/8645) and carried out with ethical approval from the Beatson Institute for Cancer Research and the University of Glasgow under the Animal (Scientific Procedures) Act 1986 and the EU directive 2010.

7-week-old CD1-nude male mice were obtained from Charles River (UK). 5 x 10^6^ PC3 cells stably expressing either Scr (12 mice) or IQSEC1 KD4 (10 mice) shRNA were surgically implanted into one of the anterior prostate lobes of each mouse ^67^. The mice were continually assessed for signs of tumour development and humanely sacrificed at an 8-week timepoint when tumour burden became restrictive. Primary tumours were imaged and tumour area and volume measured using VevoLAB ultrasound equipment and software. Significance was calculated using Mann-Whitney, *p≤0.05. Percentage of mice with tumours or with proximal (lumber lymph nodes and epididymal fat pads) or distal (thoracic lymph nodes and lungs) metastases were calculated by gross observation. p values were determined using a Chi-squared test (with fisher’s exact test) adjustment, n.s. = not significant, *p≤0.05.

### Immunohistochemistry (IHC)

4µm formalin fixed paraffin embedded (FFPE) sections were cut from tissue blocks and maintained at 60 °C for 2 hours. FFPE sections were stained with Cleaved Caspase 3 (CST, 9661) and Ki67 (Abcam, ab16667) antibodies using the Leica Bond Rx autostainer. Sections were loaded onto the autostainer, de-waxed (Leica, AR9222) and epitope retrieval carried out. Both antibodies were retrieved using ER2 buffer (Leica, AR640) for 30 minutes at 95°C. Sections were rinsed with Leica wash buffer (Leica, AR9590) before peroxidase block was performed using an Intense R kit (Leica, DS9263). Caspase 3 was stained at a previously optimised dilution of 1/500 and Ki67 at 1/100 before washing with Leica wash buffer and then application of rabbit envision secondary (Agligent, K4003) and visualized using DAB in the Intense R kit. Sections were counterstained with Hameatoxylin and mounted using DPX (CellPath, SEA-1300-00A).

IHC slides were scanned using a Leica SCN 400 F scanner at X20 magnification and images uploaded to Halo Image analysis platform (Indica Labs.). Images were analysed using the CytoNuclear v1.5 algorithm. The percentage of Ki67 or Cleaved Caspase 3 positive cells in prostate tumours or lymph nodes was calculated. Additional measurements such as the average cell area and average cytoplasmic area were also taken.

### Met uptake assay

Met uptake assays were performed using a fluorescently labelled Met-647 antibody (BD Biosciences) to assess the cells’ ability to internalise and traffic Met. Cells plated on 5 x 96 well plates were equilibrated at 37°C for 1 hour in antibody binding medium (RPMI + 0.5% BSA + 1 M CaCl_2_, pH 7.4). Cells were washed twice with ice-cold PBS, incubated with Met-647 conjugated antibody for 1 hour at 4°C (1:100) then washed again with PBS. One plate was fixed as described previously. Two plates were treated with HGF (50ng/μl) and chloroquine (100μM) diluted in pre-warmed medium for 10 or 30 minutes at 37°C. They were then acid stripped (washed three times at 4°C with 0.5M acetic acid, 0.5M NaCl in PBS) and fixed as previously described. The remaining two plates were incubated at 17°C with pre-warmed medium supplemented with HGF and chloroquine for 1 hour. Acid stripping was performed and plates maintained at 37°C without HGF or chloroquine for 10 min or 30 minutes prior to fixation. Cells were stained with Hoechst and phalloidin and imaged using the Opera Phenix™ High Content analysis system. Data are presented as line graphs that show normalised (Z-score) relative region intensities compared to the intensity of the whole cell. Box and whiskers plot shows total normalised (Z-score) intensity throughout the cells. n=2 with between 100-500 cells examined per experiment. P values were calculated using Welch’s t-test and are shown on each graph as follows; n.s. = not significant, **p≤ 0.01, ***p ≤0.001 and ****p< 0.0001.

### Protein Purification

pET20b-6xHis-ARF5D17 or pGEX-5x-1-hGGA3 [VHS-GAT] were transformed into BL21 (DE3) pLysS bacteria (Promega). Single colonies were grown in lysogeny broth (LB) medium in appropriate antibiotics until OD_600nm_ of 0.3. Protein expression was induced using 200µM isopropyl β-D-1-thiogalactopyranoside (IPTG, Sigma). Bacterial cells were pelleted and resuspended in 20ml Buffer A (for His-tagged: 20mM Tris, 300mM NaCl, 5mM MgCl2, 150µ l BME; for GST-tagged: 20mM Tris, 150mM NaCl, 5mM MgCl2, 150µl BME) containing protease inhibitors. DNAse (Sigma) was added to the lysate. Bacteria were lysed by passing through a 20,000 psi-pressurised microfluidizer. Lysates were collected at 20,000 x *g* for 40 min at 4°C and soluble proteins in the supernatant were sterile filtered 5µm PVDF Membrane (Merck).

PCR products encoding human IQSEC1-SEC7-PH (residues 517-866 based on variant 2 numbering) WT and GEF-dead (E620K) were cloned into pEGFP-C1 fused with an N-terminal 12xHis tag followed by a 6x Glycine linker replacing the GFP sequence. HEK293 suspension cells were transfected with constructs using polyethylenimine (PEI, Sigma). HEK293 suspension cells were cultured at 37°C and 8% CO_2_ and 120 rpm. Cells were collected after 4 days, resuspended in His-IQSEC1 Buffer A (40mM Tris 7.5 pH, 300mM NaCl, 5mM MgCl_2,_ 60µl *β*-mercaptoethanol, 25mM Imidozol) and sonicated. Soluble fraction was collected at 40 min, 21,000 x *g* at 4°C and sterile-filtered using a 0.45µm PVDF-membrane (Merck).

Soluble fractions were loaded onto an ÄKTA purification system (GE Healthcare) using an equilibrated HisTrap or GSTrap (both GE Healthcare) column. The protein containing fraction was loaded onto the column at 1ml/min. Proteins were eluted from columns using an imidozol gradient buffer (25–300mM) or glutathione elution buffer (150mM NaCl. 25mM Tris, 20mM Glutathion) for His-tagged or GST-tagged proteins, respectively. Protein containing fractions were concentrated (Amigon Ultra-15 centrifuge filter) and gel purified on an appropriate size exclusion column (Superose-6 10-300 GL 24ml for His-tagged IQSEC1, S75 16-600 120ml for His-tagged ARF5, S200 16-600 120ml for GST-GGA3) in Gel Filtration (GF) buffer (20mM Tris, 150mM NaCl, 1mM DTT, 5mM MgCl_2_). Fractions containing purified proteins were concentrated, snap frozen and stored at −80°C.

### Fluorescent Polarisation Assay

Nucleotide exchange of ARF5 protein was examined by observing changes in fluorescent polarisation. 100μM recombinant ARF protein was incubated with 200μM mantGDP and 50mM EDTA in Gel Filtration Buffer (GF) (20mM Tris, 150mM NaCl, 1mm DTT) overnight at 18°C. 100mM MgCl_2_ was added to stop the exchange reaction. A PD10 (GE Healthcare) desalting column was then washed with GF buffer containing 5mM MgCl_2,_ 500μl of sample added and then eluted with GF buffer. Protein concentration was determined using a Bradford assay as per manufacturers instructions. 20μM of GDP nucleotide and 2μM of GEF protein was sequentially added to 1μM ARF and polarisation changes measured with a Photon Multiplier Detection System (Photon International Technology). Excitation was set to 366 nm and emission to 450 nm. Inhibitors were added to the reaction prior to addition of GEF protein.

### Proliferation Assays

PC3 cells were plated in a 96-well ImageLock plate, in triplicate for 24 hours. Imaging was carried out on IncuCyte® ZOOM each hour for 48 hours. Cell area per well was then measured using the IncuCyte® ZOOM analysis software. n=3 independent experiments with 3 replicates per condition. P-values (Student’s t-test): **p≤0.01 and ****p≤0.0001.

### Invasion and Migration Assays

ImageLock plates were coated with 10% GFRM diluted in medium overnight at 37°C. Cells were re-suspended in 100μl medium and plated for 4 hours at 37°C. The resultant monolayer was wounded using a wound making tool (Essen Biosciences), washed twice with medium to remove debris, and overlaid with either 50μl of 25% GFRM for invasion assays or 100μl medium for migration assays. After an hour at 37°C 100μl medium was added to invasion assays and plates imaged every hour for 4 days using the IncuCyte® ZOOM. Sample number (n) and replicate number are stated in the appropriate Figure Legend. Results are presented as Relative Wound Density (RWD) for each time point and for the time point at which the average RWD of the control samples is 50% (Tmax1/2). Values are mean ± s.d. P values were calculated using Student t-Test and are shown on each graph as follows; n.s. = not significant, *p≤0.05, **p≤ 0.01, ***p ≤0.001 and ****p< 0.0001.

### Immunoblotting

Cells were plated for 24 hours prior to treatment with growth factors or inhibitors for a further 24 hours. Plates were washed twice with ice cold PBS then Lysis Buffer added for 15 minutes (50mM Tris-HCl, pH 7.4, 150mM NaCl, 0.5mM MgCl_2_, 0.2mM EGTA, and 1% Triton X-100 with cOmplete protease inhibitor cocktail and PhosSTOP tablets (Roche). Cells were scraped and lysates clarified by centrifugation at 13,000 rpm at 4°C for 15 minutes. BCA Protein Assay kit (Pierce) was used to determine protein concentration. SDS-PAGE was then performed and proteins transferred to PVDF membranes using the iBlot 2 transfer system (Thermo Fisher Scientific). Membranes were incubated for an hour in Rockland blocking buffer (Rockland) and primary antibodies added overnight at 4°C (1:1000 unless stated otherwise). Antibodies used were as follows: anti-GAPDH (CST 2118 1:5000), anti-IQSEC1 (Sigma G4798), anti-IQSEC1 (Caltag-Medsystems PSI-8009), anti-GFP (Merck 000000011814460001), anti-LRP1 (Sigma L2295), anti-Met (CST 3127), anti-Met phospho 1234/1235 (CST 3077), anti-Akt (CST 2920), anti-Akt phospho S473 (CST 3787), anti-ARF1 (Novus Biologicals NB-110-85530), anti-ARF6 (Sigma A5230), ARF5 (Novus Biologicals H00000381-M01), anti-ARFGAP1 (Sigma HPA051019), anti-SORL1 (BD 611860), anti-Sin1 (CST 12860), anti-RICTOR (CST 2114), anti-PPC6 (Sigma HPA050940 1:250), anti-14-3-3ζ/Δ (CST 7413), anti-PIP5Ka (CST 9693) and PIP5Kb (Sigma K0767). After addition of appropriate secondary antibodies for 45 minutes, membranes were washed three times in TBST and imaged using a ChemiDoc Imager (BioRad) or Odyssey Imaging System (LI-COR Biosciences). Sample number (n) is stated in the appropriate Figure Legend and quantitation is shown as mean ± s.d. P values were calculated using Student t-test and are shown on each graph as follows; n.s. = not significant, *p≤0.05, **p≤ 0.01, ***p ≤0.001 and ****p< 0.0001. GAPDH was used as a loading control for each immunoblot and a representative image for each sample set is shown where appropriate.

### GFP Trap and Immunoprecipitation

Immune complexes were collected when 1mg of cell lysate was immunoprecipitated with 2μg of anti-Met antibody (CST 3127) overnight at 4°C with rotation. Anti-mouse agarose or mouse agarose (both Sigma) were added for 1 hour at 4^0^C prior to 3 washes in Lysis Buffer. Samples were then separated by SDS–PAGE, transferred to a PVDF membrane and immunoblotted. Lysates from cells expressing GFP tagged proteins were immunoprecipitated using a GFP-Trap Kit (Chromotek) as per manufacturer’s instructions and immunoblotting performed as above. n=3 and quantitation is shown as mean ± s.d.

### Mass Spectrometry

For Mass Spectrometry analysis agarose beads were resuspended in a 2M Urea and 100mM ammonium bicarbonate buffer and stored at −20°C. On-bead digestion was performed from the supernatants. Triplicate biological replicates were digested with Lys-C (Alpha Laboratories) and trypsin (Promega) on beads as previously described^68^. Tryptic peptides were separated on a 20cm fused silica emitter (New Objective) packed in house with reverse phase Reprosil Pur Basic 1.9μm (Dr Maisch GmbH) using an EASY-nLC 1200 (Thermo Fisher Scientific) coupled online to an Orbitrap Q-Exactive HF mass spectrometer (Thermo Fisher Scientific) via nanoelectrospray ion source (Thermo Fisher Scientific). For the full scan a resolution of 60,000 at 250Th was used. The top ten most intense ions in the full MS were isolated for fragmentation with a target of 50,000 ions at a resolution of 15,000 at 250Th. MS data acquisition were performed using the XCalibur software (Thermo Fisher Scientific). The MaxQuant software version 1.5.5.1^69^ was used to process MS Raw files and searched with Andromeda search engine^70^, querying UniProt ^71^. Database was searched requiring specificity for trypsin cleavage and allowing maximum two missed cleavages. Methionine oxidation and N-terminal acetylation were specified as variable modifications, and Cysteine carbamidomethylation as fixed modification. The peptide and protein false discovery rate (FDR) was set to 1%. The common reverse and contaminant hits (as defined in MaxQuant output) were removed. Only protein groups identified with at least one uniquely assigned peptide were used for quantification. For label-free quantitation, proteins quantified in all 3 replicates in at least one group, were measured according to the label-free quantitation algorithm available in MaxQuant ^72^. Significantly enriched proteins were selected using a Welch t-test with a 5% FDR (permutation based). The mass spectrometry proteomics data have been deposited as partial submission to the ProteomeXchange Consortium via the PRIDE partner repository ^73^ with the dataset identifier PXD013810. Hits were prioritised based on fold change expression data mined from publicly available RNAseq data from PC3 sublines from published data sets (PC3E versus GS689.Li; SRS354082) and PC3-Epi versus PC3-EMT14; GSE48230 ^40, 41^.

### GGA3 Pulldown

A pulldown assay was performed to examine GTP-loading of ARF proteins using the ARF-GTP-specific binding domain of GGA3. Cells were cultured for 48 hours then serum starved overnight. HGF in serum free medium was added for 30 minutes. Cells were then lysed on ice in pulldown-lysis buffer (50mM Tris, 100mM NaCl, 2mM MgCl2, 0.1% SDS, 0.5% Na-deoxycholate, 1% Triton X-100, 10% glycerol), syringed 5 times and centrifuged at 4°C 14,000*g* for 1 minute. Spin columns were equilibrated with 50μl of Glutathione Agarose resin and washed with pulldown-column wash buffer (1:1 pulldown-lysis buffer and 1xTBS). 80μg of GST-GGA3-GAT recombinant fusion protein was immobilised on the agarose resin by incubation at 4°C with gentle rocking for 1 hour. 1.5mg of lysate was added onto spin columns and incubated at 4°C for 2 hours with rocking. Unbound proteins were washed off the column with pulldown-wash buffer (50mM Tris, 100mM NaCl, 2mM MgCl2, 1% NP-40, 10% glycerol). 60μl of pulldown-elution buffer (10mM Glutathione in 1xTBS) was added to the spin column and incubated for 5 min at RT. Eluted protein was collected at 1,250*g* for 1 minute and samples were prepared for SDS-PAGE and western blotting as described above.

### qPCR

RNA was extracted using an RNeasy kit (Qiagen) and reverse transcription of RNA performed using a High-Capacity cDNA Reverse Transcription Kit (Thermo Fisher Scientific) following the manufacturer’s protocols. TaqMan qPCR was then carried out as per manufacturer’s instructions (Thermo Fisher Scientific). n = 3 technical replicates. Data were analysed using the Applied Biosystems 7500 Software v2.0.6 and the relative quantitation (RQ) was calculated using the comparative C_t_ (ΔΔC_t_) method. *ARF* and *ARFGEF* expression were detected using a custom qPCR primer panel. Details are available on request.

### Analysis of patient cohorts

The majority of patient data (Copy number, mutational status, RNAseq, RPPA, co-expression matrices and clinical annotation) was accessed, analysed and downloaded using in-platform cBioportal.org tools ^74, 75^. Normal versus Tumour RNAseq, including transcript variant expression, was obtained by downloading IQSEC1 variant annotation across the TCGA pan cancer dataset using the TCGA Splicing Variants Database (www.TSVdb.com) ^76^. RPPA and IQSEC1 variant data was analysed using custom KNIME Analytics platform pipelines.

## Supporting information

Graphical Abstract

## Acknowledgements

This work was supported by the following grants; D.M.B. NIH K99CA163535, M.N. CRUK (C596/A19481), K.N. CRUK (C7932/A25170), E.S., AR.-F, L.M. S.Z., and S.L. CRUK C596/A17196, E.M. CRUK A25142, S.I. and T.Y. CRUK (A19257). We would like to thank the Core Services and Advanced Technologies at the Cancer Research UK Beatson Institute, with particular thanks to the Beatson Advanced Imaging Resource, Histology and Molecular Technologies.

## Author Contributions

M.N., D.M.B., K.N., E.S. designed experiments and analysed data. M.N., D.M.B., K.N., E.S., T.Y., A.R-F, R.P., L.G. and S.M. performed experiments and analysis. L.M. and E. Shanks helped with the development of high-throughput imaging and analysis. S.L. and S.Z. performed and analysed the proteomic experiments. T.Y. and S.I. guided protein purification and biochemistry. D.M.B. and E.M. analysed patient data. S.M., R.P., and L.G. performed and analysed, while K.B. and H.L. guided i*n vivo* experiments. J.P.M. generated and/or provided cell lines. D.M.B, M.N. and E.S. wrote the manuscript. D.M.B. supervised the study. All authors discussed the study and commented on the manuscript.

---

**Supplementary Figure 1. IQSEC1 is a regulator of cell elongation and collective invasion.**

**(A)** Western blot of PC3 cells stably expressing Scrambled (Scr) or *IQSEC1* (KD1, KD4) shRNAs using anti-IQSEC1 and GAPDH (loading control for IQSEC1 blot) antibodies. Arrowheads indicate reduction of three major bands. IQSEC1 intensity for all bands combined normalised to Scr control is shown. n=3 independent experiments. Values are mean ± s.d. p-values (Students t-test): *p≤0.05, **p≤0.01.

**(B)** PC3 cells expressing Scr or *IQSEC1* KD shRNA were imaged every hour for 3.5 days. Cell confluence was measured and normalised to control. Values are mean ± s.d. p-values (Students t-test) are annotated in plot, **p≤0.01 and ***p≤0.001.

**(C)** Schematic of phenotypic analysis of PC3 cells in 2D. Machine learning applied to confocal images to classify and quantify cells into three categories based on shape: spindle, spread, round.

**(D)** PC3 cells expressing Scr or *IQSEC1* KD4 shRNA were stained with whole cell stain (WCS) (red) and Hoechst/nuclei (blue) (left panels). Cells were classified into spindle, spread and round phenotypes shown in green, red and blue respectively (right panels). Scale bar, 100μm.

**(E-F)** Proportion of PC3 cells with each phenotype **(E)** or classified into round, spindle or spread **(F)** is shown. Heatmap indicates log2 fold change of each phenotype over Scr (upper heatmap). p-values (one-way ANOVA): greyscale values as indicated (lower heatmap). n=4 independent experiments, 10 replicates per condition, minimum 118,000 cells imaged/condition in total.

**(G-H)** Schema, PC3 cells plated for 24 hours before the resultant monolayers were **(G)** wounded or **(H)** wounded and overlaid with 25% ECM for 1 hour prior to imaging using 10x objective. Phase contrast images are shown where the yellow dashed lines indicate initial scratch wound and red pseudo colour shows wound after 24 hours. Magnified images of boxed areas are shown for different time points. Relative wound density (RWD) was calculated at each time point and graphed (lower left panels). RWD at the time point where the Scr controls are 50% closed (t=Max_1/2_) is also shown (lower right panels). Samples were normalised to the average of all Scr controls across experiments. n=3 independent experiments, 3 replicates/experiment in **G** and 4 in **H**. Values, mean ± s.d. p-values (Student’s t-test): *p≤0.05, **p≤0.01 and ****p≤0.0001.

**(I)** Schema, depicts invasive ability of PC3 in ECM in the presence or absence of IQSEC1.

**(J)** Schema, summarizes the effect IQSEC1 depletion has on 2D growth, shape and migration and on 3D invasion.

**Supplementary Figure 2. IQSEC1 v2 induces spindle shape and protrusive invasion.**

**(A)** Schema, outlines generated by CellProfiler were colour-coded at each time point and overlaid in 12 hour blocks using Fiji. Scale bars, 20μm.

**(B)** Phase contrast images are shown for PC3 acini expressing Scr or *IQSEC1* KD4 shRNA. Acini outlines, yellow and protrusions, white arrowheads. Magnified images of boxed regions are shown at different time points. Scale bars, 100μm.

**(C)** Quantitation of images shown in **B**. CellProfiler was used to measure invasion parameters i.e. area and compactness, over 4 days, in 12 hour blocks. Quantitation shown as Z-score-normalised values (upper heatmap). p-values (one-way ANOVA): greyscale values as indicated (lower heatmap). n=4 independent experiments, 4 replicates/condition, 1,555-2,035 acini/condition.

**(D)** 2D PC3 cells expressing GFP or GFP-IQSEC1 v1-4 and either Scr or *IQSEC1* KD4 shRNA classified into spindle, spread and round phenotypes shown in green, red and blue respectively (right panels). Scale bars, 100μm. Quantitation is shown in heatmaps as log2 fold change of each phenotype over control (top row in left heatmap). p-values (one-way ANOVA): greyscale values as indicated (right heatmap). n=2 independent experiments, 4 replicates/condition, 1,431-5,587 cells quantified/condition in total.

**(E-H)** PC3 acini expressing GFP or GFP-IQSEC1 v2 or v4 and either Scr or *IQSEC1* KD4 shRNA were analysed for **(E)** Nuclei number, **(F)** Acinus volume (μm^3^), **(G)** proliferation (Ki67) or **(H)** apoptosis (CC3, Cleaved Caspase 3). In **(F-H)** values, log2-normalised fold change to average. p-values (one-way ANOVA): **p≤0.01, ***p≤0.001 and ****p≤0.0001. n = 2 independent experiments with 4 replicates/condition. Between 365 – 724 cells quantified per condition in total.

**(I)** 2D PC3 cells expressing GFP or GFP-IQSEC1 v1-4 and either Scr or *IQSEC1* KD4 shRNA were also analysed for nuclear, cytoplasmic and cortical localisation of GFP using Harmony High-Content Imaging and Analysis Software. Quantitation is shown in heatmaps. n=2, 4 replicates/condition.

**(J)** Schema, depicts how optical slices were combined to form maximum projections of acini body and protrusions.

**(K)** Maximum projections of PC3 acini fixed and stained with F-actin (black) and either an IQSEC1 antibody that detects all variants (upper panels) or one that is specific for v2 (lower panels). Localisation of IQSEC1 can be appreciated from FIRE LUT in magnified images of protrusions. Scale bars, 10μm.

**Supplementary Figure 3. N-terminal common extension in IQSEC1 v2 confers enhanced invasive activity in a GEF-dependent manner.**

**(A)** Schemes, IQSEC1 chimeras generated by swapping N- and C-terminal domains from IQSEC1 variants v1-v4. Colour coding as follows: v1 = blue, v2 = pink and v4 = green. Chimeras with alternate N-termini, C-termini or are GEF dead are labelled (a), (b) or (c) respectively.

**(B)** PC3 cells expressing GFP, GFP-IQSEC1 v2 or GFP-IQSEC1 chimeras and either Scr or *IQSEC1* KD4 shRNA were classified into spindle, spread and round phenotypes. n=2 independent experiments, 4 replicates/condition, 10,000-35,000 cells/condition in total.

**(C)** Cells from **B** were also analysed for nuclear, cytoplasmic and cortical localisation and quantitation is shown in heatmaps. n=2 independent experiments, 4 replicates/condition.

**(D)** PC3 acini expressing GFP, GFP-IQSEC1 v2 or GFP-IQSEC1 chimeras were fixed after 3 days and FIRE LUT used to show localisation and intensity of GFP. Magnified images of boxed regions are also shown in lower panels. Scale bars, 10μm.

**(E)** Phase images from PC3 acini expressing GFP, GFP-IQSEC1 v2 or GFP-IQSEC1 chimeras and either Scr or *IQSEC1* KD4 shRNA are shown after 96 hours. GFP-positive acini are outlined in yellow. Scale bars, 100μm.

**(F)** Quantitation of images shown in **E**. n=2 independent experiments, 4 replicates/condition, 550-1,662 acini/condition in total. Cartoon, depicts acini phenotype representative of each condition. **(G)** Schema, depicts the role of IQSEC1 v1 and v2 termini on localisation and invasion.

**Supplementary Figure 4. ARF5 and ARF6 co-operate to regulate both growth and invasion.**

**(A)** Quantitation of shape classification of PC3 cells expressing Scr, *ARF5, ARF6* or *AR5/6* shRNA is shown. Heatmaps show log2 fold change of each phenotype over control (left heatmaps). p-values (one-way ANOVA): greyscale values as indicated (right heatmaps). n=2 independent experiments, 4 replicates. Minimum of 4,000 cells/condition analysed in total.

**(B)** Schema, comparison of activation of WT ARF GTP and fast-cycling ARF mutants.

**(C)** Western blot of PC3 cells expressing mNeon Green (mNG), ARF5-mNG, ARF5- mNG fast cycling (TA), ARF6-mNG or ARF6-mNG fast cycling (TN) using anti-ARF5, ARF6 and GAPDH (loading control for ARF5 blot) antibodies.

**(D)** Quantitation of shape classification of PC3 cells described in **C** is shown. n=2, 4 replicates.

**(E)** Quantitation of PC3 acini described in **C**. Heatmaps show area and compactness measurements as Z-score-normalised values (upper heatmap). p-values (one-way ANOVA): greyscale values as indicated (lower heatmap). n=2 independent experiments with 4 replicates/condition. Minimum of 900 acini/condition analysed in total. Cartoon, depicts acini phenotype representative of each condition.

**(F)** Quantitation of shape classification of PC3 cells expressing mNG, ARF5-mNG and ARF6-RFP or ARF5-mNG TA and ARF6-RFP TN is shown. n=3 independent experiments, 3 replicates.

**(G)** Quantitation of PC3 acini. n=2 independent experiments, 4 replicates. Cartoon, depicts acini phenotype representative of each condition.

**(H)** Quantitation of shape classification of PC3 cells expressing control or ARF5/ARF6 and Scr or *IQSEC1* KD4 shRNA. n=2 independent experiments, 4 replicates.

**(I)** PC3 cells expressing Scr or *IQSEC1* shRNA were serum starved overnight and HGF was then added for 30 minutes. GGA3 pulldown was carried out and western blot analysis performed using anti-IQSEC1, ARF5, ARF6, and GAPDH (loading control for ARF6 blot) antibodies. Quantitation of ARF5 and ARF6 GTP loading normalised to control is shown (right panels). n=3 independent experiments. Values, mean ± s.d. p-values (one-way ANOVA): no significance.

**(J)** Schema, modulation of ARF activity using SecinH3, NAV-2729 or QS11.

**(K)** *In vitro* fluorescent polarisation assay was carried out to study exchange of fluorescent nucleotides (MANT) on ARF5. IQSEC1 v2 SEC7-PH WT and GEF dead mutant IQSEC1 v2 SEC7-PH GEF* were used to test the role of IQSEC1 in activation of ARF5. ARFs, nucleotides, GEFs and inhibitors were added at time points indicated. Changes in polarisation over time are shown.

**(L)** Western blot of PC3 cells treated for 24 hours with NAV-2729, SecinH3 or QS11 using anti-IQSEC1, ARF5, ARF6 and GAPDH (loading control for IQSEC1) antibodies.

**(M)** Representative phase contrast images of PC3 acini treated with NAV-2729, SecinH3 or QS11 after 96 hours. Scale bars, 100μm.

**(N)** Quantitation of PC3 acini shown in **M**. n=1 experiment, 4 replicates.

**(O)** Cartoon, depicts representative acini phenotype upon treatment with inhibitors.

**Supplementary Figure 5. IQSEC1 interacts with multiple transmembrane proteins.**

**(A)** GFP-trap immunoprecipitation was performed on PC3 cells expressing GFP, GFP-IQSEC1 v2 or v4. Western blot analysis was then carried out using anti-IQSEC1, LRP1, Met, SORL1, RICTOR, Sin1, ARFGAP1, 14-3-3ζ/Δ, PPC6, GFP and GAPDH (loading control for GFP blot) antibodies.

**(B)** Western blot of PC3 subclones using anti-IQSEC1, ARF5, ARF6, ARFGAP1, LRP1, phospho-Y1234/1235 Met, Met, SORL1, RICTOR and GAPDH (loading control for IQSEC1 blot) antibodies.

**(C)** Western blot of Epi and EMT14 PC3 subclones using anti-IQSEC1, ARF5, ARF6, ARFGAP1, LRP1, phospho-Y1234/1235 Met, Met, SORL1, RICTOR and GAPDH (loading control for IQSEC1 blot) antibodies.

**(D)** Met immunoprecipitation was carried out on PC3 cells expressing Scr or *IQSEC1* KD4 shRNA. Western blotting was then performed on lysates (upper panels) and IPs (lower panels) using anti-LRP1, Met, IQSEC1 and GAPDH (loading control for IQSEC1 blot) antibodies. Quantitation of LRP1/Met interaction normalised to Scr is shown. Values, mean ± s.d. n=3 independent experiments. p values (one-way ANOVA): n.s. not significant.

**(E-F)** Western blot analysis of PC3 cells expressing Scr or **(E)** *LRP1* and **(F)** *Met* shRNA. GAPDH is the loading control for the Akt blots.

**(G)** Quantitation of shape classification of PC3 cells expressing shRNAs for IQSEC1 binding partners as log2 fold change of each phenotype over control (left heatmap). p-values (one-way ANOVA): greyscale values as indicated (right heatmap). n=2, 4 replicates per condition, minimum 14,000 cells imaged/condition.

**(H)** Quantitation of PC3 acini expressing GFP or LRP1-GFP. Heatmaps for area and compactness measurements shown as Z-score-normalised values (upper heatmap). p-values (one-way ANOVA): greyscale values as indicated (lower heatmap). n=3 independent experiments, 3 replicates, minimum 150 acini/condition in total.

**(I)** Western blot of PC3 cells expressing *SORL1* shRNA. GAPDH is the loading control for the Akt blots.

**(J)** Representative phase contrast images of PC3 acini expressing Scr (also shown in **4H**), *SORL1_*2 and *RICTOR_2* shRNA taken after 96 hours. Scale bars, 100μm.

**(K)** Quantitation of PC3 acini shown in **J**. n=2 independent experiments, 4 replicates, minimum 700 acini/condition in total.

**(L-N)** Western blots of PC3 cells expressing **(L)** *RICTOR* or **(M)** *SIN1* or **(N)** *ARFGAP1* shRNA. GAPDH is the loading control for Akt blots.

**(O)** Representative phase contrast images of PC3 acini expressing Scr, *ARFGAP1_1* and *ARFGAP1_2* shRNA taken after 96 hours. Scale bars, 100μm.

**(P)** Quantitation of PC3 acini shown in **O**. n=2 independent experiments, 4 replicates, minimum 1,100 acini/condition in total.

**(Q)** Cartoon, depicts representative acini phenotype upon depletion of ARFGAP1 and ARF GTPase cycle.

**Supplementary Figure 6. HGF stimulation is abrogated by IQSEC1 depletion.**

**(A)** Western blot of PC3 cells expressing Scr or *IQSEC1* KD4 shRNA stimulated with HGF for 30 minutes using anti-IQSEC1, phospho-Y1234/1235 Met, Met, phospho-S473 Akt, Akt, ARF5, ARF6 and GAPDH (loading control for Akt blot and sample control for ARFs) antibodies.

**(B)** Quantitation of Met and Akt: phospho/total, phospho/GAPDH and total/GAPDH. Expression is presented as signal intensity relative to control. Values, mean ± s.d. n=3 independent experiments. p-values (Student’s t-test): *p≤0.05, **p≤0.01, ***p≤0.001 and ****p≤0.0001.

**(C)** Phase contrast images of PC3 acini expressing Scr or *IQSEC1* KD4 shRNA stimulated with HGF for 96 hours. Scale bars, 100μm. Cartoon, depicts acini phenotype representative of each condition.

**(D)** Quantitation of PC3 acini shown in **C**. Heatmaps show area and compactness measurements as Z-score-normalised values (upper panels). p-values (one-way ANOVA): greyscale values as indicated (lower heatmap). n=3 independent experiments, 4 replicates/condition, 1,400 - 4,025 acini/condition in total.

**(E)** Western blots of PC3 cells expressing Scr or *IQSEC1* KD4 shRNA treated with cycloheximide (CHX) for various times using anti-IQSEC1, Met and GAPDH (shown for Met blot) antibodies. Quantitation of Met expression levels normalised to time 0 is shown. Values, mean ± s.e. n=3 independent experiments. p-values (one-way ANOVA): *p≤0.05, ***p≤0.001 and ****p≤0.0001.

**Supplementary Figure 7. IQSEC1-ARF controls cortical PI(4,5)P_2_ generation, which is required for production of PIP*_3_*.**

**(A)** PC3 cells expressing mNeonGreen tagged PH-PLCδ (PIP_2_) or PH-Grp1 (PIP_3_) and either Scr or *IQSEC1* shRNA KD4 were fixed after 2 days. FIRE LUT is used to show localisation and intensity of GFP in magnified images of boxed regions. White arrowhead indicates localisation at protrusive tip. Scale bars, 20µm.

**(B-C)** Quantitation of **(B)** PI(4,5)P_2_ or **(C)** PIP_3_ cortical enrichment in spindle, spread and round cells expressing either Scr or IQSEC1 KD4 shRNA is shown in box-and-whiskers plot. n=3 independent experiments, 4 replicates/condition/experiment. 1,330/1,049 and 2,222/1,182 cells analysed in total for Src/*IQSEC1* KD4 shRNA respectively in **B** and **C**. p-values (one-way ANOVA).

**(D)** Cartoon, summary of cortical PI(4,5)P_2_ and PIP_3_ localization in different cell shapes.

**(E)** PC3 cells were fixed and stained with anti-PI(4,5)P_2,_ PI(3,4,5)P_3_ antibodies, F-actin (green outlines) and Hoechst (nuclei, magenta). Magnified images and FIRE LUT of boxed regions are shown. Arrowheads indicate peripheral localization of PIP antibodies. Scale bars, 10µm.

**(F-G)** Quantitation of the total intensity/cell of anti-PI(4,5)P_2,_(left panels) or PI(3,4,5)P_3_ (right panels) antibodies in cells treated with LY294002 **(F)** or over-expressing PI3K-FLAG **(G)** is shown in box-and-whiskers plot. n=1 experiment, **(F)**, 2 **(G)** 4 replicates/condition, 680 – 1,410 **(F)** and 2,391 – 3,826 (FLAG +) **(G)** cells/condition. p-values (Students t-test).

**(H)** PC3 cells expressing mNeonGreen tagged PH-Grp1 (PIP_3_) in the absence or presence of *IQSEC1* were treated with LY294002 overnight, fixed and stained with F-actin, CellMask and Hoechst. Quantitation of peripheral/total PIP_3_ intensity/cell is shown in box-and-whiskers plot. n=2 independent experiments, 4 replicates/condition, 220 - 241 cells/condition in total. p-values (one-way ANOVA).

**(I)** Quantitation of the total intensity/cell of anti-PI(4,5)P_2,_(left panel) or PI(3,4,5)P_3_ (right panel) antibodies in the absence or presence of *IQSEC1* is shown in box-and-whiskers plot. n=3 independent experiments, 3 replicates/condition, 470 - 499 cells/condition in total. p-values (Students t-test).

**(J)** PC3 cells expressing mNeonGreen tagged PH-Grp1 (PIP_3_) in the absence or presence of *IQSEC1* were fixed and stained with F-actin, CellMask and Hoechst. Harmony High-Content Imaging and Analysis Software was used to segment the cells into peripheral, cytoplasmic and nuclear regions using either F-actin or CellMask to define peripheral region. Quantitation of peripheral/total PIP_3_ intensity/cell is shown in box-and-whiskers plot. n=1 experiment, 4 replicates/condition, > 250 cells/condition in total. p-values (one-way ANOVA).

**(K)** Quantitation of the total intensity/cell of anti-PI(4,5)P_2,_(left panel) or PI(3,4,5)P_3_ (right panel) antibodies in cells expressing GFP, GFP-IQSEC1 v2 or GFP-IQSEC1 v4 and either Scr or *IQSEC1* KD4 shRNA is shown in box-and-whiskers plot. n=2 independent experiments, 4 replicates/condition, 511 - 1,677 GFP + cells/condition in total. p-values (Students t-test).

All box-and-whiskers plot: 10–90 percentile; +, mean; dots, outliers; midline, median; boundaries, quartiles. All p-values: n.s. not significant, *p≤0.05, **p≤0.01, ***p≤0.001 and ****p≤0.0001.

**Supplementary Figure 8. PIP5K1β-PI3Kβ-Akt pathway involved in 3D growth and invasion.**

**(A)** Western blot of PC3 cells treated with LY294002 (pan PI3K), AZD8835 (PI3Kα), AZD8186 (PI3Kβ), AS605240 (PI3Kγ), Cal-101 (PI3Kδ) and AktII (Akt) inhibitors for 24 hours. Anti-phospho-S473 Akt, Akt and GAPDH antibodies were used.

**(B)** Phase contrast images of PC3 acini described in **A**. Scale bars, 100μm.

**(C)** Quantitation of PC3 acini shown in **B**. Heatmaps show area and compactness measurements as Z-score-normalised values (upper heatmaps). p-values (one-way ANOVA): greyscale values as indicated (lower heatmaps). n=2 independent experiments, 4 replicates, minimum of 350 acini/condition in total.

**(D)** Cartoon, depicts acini phenotype representative of each condition.

**(E)** Western blot of PC3 cells expressing Scr, PIP5K1α or PIP5K1*β* shRNA using anti-PIP5K, IQSEC1, phospho-S473 Akt, Akt and GAPDH (loading control for Akt blots) antibodies.

**(F)** Phase contrast images of PC3 acini expressing Scr, PIP5K1α or PIP5K1*β* shRNA at 96 hours. Scale bars, 100μm.

**(G)** Quantitation of PC3 acini shown in **F**. n=2 independent experiments, 4 replicates, minimum of 1,348 acini/condition in total. Cartoon, depicts acini phenotype representative of each condition.

**(H)** PC3 cells expressing Scr or *IQSEC1* KD4 shRNA were stained for pS473 Akt (black) and Hoechst /nuclei (magenta). Blue arrowheads indicate localisation of active Akt. Scale bars, 10μm.

**(I-J)** Quantitation of pAkt **(I)** mean cortical/total intensity/cell or **(J)** mean cortical/total spot area/cell of PC3 cells expressing either Scr or IQSEC1 KD4 shRNA. Box-and-whiskers plots: 10–90 percentile; +, mean; dots, outliers; midline, median; boundaries, quartiles. n=3 independent experiments, 4 replicates/condition, 2,765 – 2,837 cells/condition in total. p values (one-way ANOVA): ****p≤0.0001.

**(K)** Cartoon, depicting phospho-Akt intensity, in the presence or absence of IQSEC1, at different subcellular locations.

**Supplementary Figure 9. IQSEC1 regulates growth and invasion in multiple prostate cancer cell lines.**

**(A)** Heatmap shows mRNA expression levels, mined from CCLE, of the Met-PI3K-Akt pathway across different prostate cancer cell lines.

**(B)** Western blot of prostate cancer cell lines using anti-IQSEC1, phospho-Y1234/1234 Met, total Met, LRP1, ARF5, ARF6 and GAPDH antibodies. GAPDH is a loading control for ARF6 blot.

**(C)** Schema, effect of NAV-2729 on ARF GTPase cycle.

**(D)** Phase images of acini (− and + HGF) at different time points are shown. Acini were also treated with NAV-2729 for 96 hours. Scale bars, 100μm.

**(E)** Schema, summarizes the effect of each treatment described in **D** on acini growth and invasion.

**(F-I)** Quantitation of 22Rv1 **(F)**, LNCaP **(G)**, DU145 **(H)** and DU145 + HGF **(I)** acini formation in the absence or presence of NAV-2729. 22Rv1 and DU145 acini were also expressing LRP1-GFP and IQSEC1-GFP v2 respectively. Heatmaps show area and compactness measurements as Z-score-normalised values (upper panels). p-values (one-way ANOVA): greyscale values as indicated (lower heatmap). n=2 independent experiments, 3 replicates/condition, 400 - 1,600 acini/condition in total.

**Supplementary Figure 10. IQSEC1 regulates growth and invasion in multiple murine and human cancer cell lines.**

**(A)** Phase images of acini expressing either Scr or *IQSEC1* KD4 shRNA in DU145 (− and + HGF), MDA-MB-231, KC Pten^fl/+^ or TKCC-07 are shown. Scale bars, 100μm.

**(B-E)** Quantitation of DU145 (− and + HGF) **(B)**, MDA-MB-231 **(C)**, KC Pten^fl/+^ **(D)** and TKCC-07 **(E)** acini formation in the absence or presence of *IQSEC1* KD4 shRNA. Heatmaps show area and compactness measurements as Z-score-normalised values (upper panels). p-values (one-way ANOVA): greyscale values as indicated (lower heatmap). n=3 independent experiments, 4 replicates/condition, 5,883 - 15,437 **(B)**, 4,937 - 8,114 **(C)**, 5,003 – 7,450 **(D)** and 2,608 – 5,211 **(E)** acini/condition in total.

**(F)** Schema, summarizes the effect of IQSEC1 loss on acini growth and invasion.

**Supplementary Figure 11. Additional characterisation of IQSEC1 association with clinical metrics.**

**(A)** Co-occurrence bubble plot matrix for IQSEC1, ARF5 and ARF6 CN gain compared to the 14 most common mutations, 6 CNAs, and five gene fusion events in prostate cancer. Colouring represents Log2 of odds ratio. Red, co-occurrence; blue, mutual exclusivity. Circle size, q-value. +, CN amplification (AMP) or GAIN; -, homozygous deletion (HOMDEL) or heterozygous Loss (HETLOSS). Q-values, one-sided Fisher exact test with Benjamini-Hochberg adjustment.

**(B)** CN increase frequencies (percentage of patients/cohort) in IQSEC1, ARF5, ARF6 across cancer types from pan-cancer TCGA analyses.

**(C-D)** Differential Copy Number Alteration **(C)** or **(D)** mutation frequencies between IQSEC1 CN-amplified patients compared to non-amplified comparison group. Red, IQSEC1 CN increase-associated. Blue, control group-associated. N=9,892 patients, across 32 tumour types. Q-values, one-sided Fisher exact test with Benjamini-Hochberg adjustment.

**(E)** Bubble heatmap of protein changes related to IQSEC1 expression in patients. Values represent significant changes in protein/phospho-proteins profiled by RPPA analysis that are consistently altered in the same direction in at least a quarter of pan-cancer TCGA cohorts when patients were grouped by median split of IQSEC1 mRNA expression. Values are Log2-transformed difference between a median split of total IQSEC1 mRNA levels. Red, co-occurring with high IQSEC1; blue, associated with low IQSEC1 levels. In RPPA, 190 protein/phospho-proteins profiled. Circle size, p-value. P-values, Independent Groups T-test. Patient sample n=7,790; See Table S3 for breakdown by tumour type.

**Table S1. IQSEC1 variant nomenclature.** Reference identification numbers and nomenclature for IQSEC1 variants.

**Table S2. Normal versus tumour IQSEC1 variant and expression values and patient same sizes.** Values corresponding to Bubble Heatmap in Figure 8S, listed by tumour type.

**Table S3. Patient sample count for TCGA Pan-cancer RPPA data per tumour type.**

## References

1. Halaoui, R. & McCaffrey, L. Rewiring cell polarity signaling in cancer. Oncogene 34, 939–950 (2015).

2. Pampaloni, F., Reynaud, E.G. & Stelzer, E.H. The third dimension bridges the gap between cell culture and live tissue. Nature reviews. Molecular cell biology 8, 839–845 (2007).

3. Bryant, D.M. & Mostov, K.E. From cells to organs: building polarized tissue. Nature reviews. Molecular cell biology 9, 887–901 (2008).

4. Zajac, O. et al. Tumour spheres with inverted polarity drive the formation of peritoneal metastases in patients with hypermethylated colorectal carcinomas. Nature cell biology 20, 296–306 (2018).

5. Bryant, D.M. et al. A molecular switch for the orientation of epithelial cell polarization. Developmental cell 31, 171–187 (2014).

6. Shamir, E.R. & Ewald, A.J. Three-dimensional organotypic culture: experimental models of mammalian biology and disease. Nature reviews. Molecular cell biology 15, 647–664 (2014).

7. Parachoniak, C.A. & Park, M. Dynamics of receptor trafficking in tumorigenicity. Trends in cell biology 22, 231–240 (2012).

8. Joffre, C. et al. A direct role for Met endocytosis in tumorigenesis. Nature cell biology 13, 827–837 (2011).

9. Menard, L., Parker, P.J. & Kermorgant, S. Receptor tyrosine kinase c-Met controls the cytoskeleton from different endosomes via different pathways. Nature communications 5, 3907 (2014).

10. Parachoniak, C.A., Luo, Y., Abella, J.V., Keen, J.H. & Park, M. GGA3 functions as a switch to promote Met receptor recycling, essential for sustained ERK and cell migration. Developmental cell 20, 751–763 (2011).

11. Humphreys, D., Davidson, A.C., Hume, P.J., Makin, L.E. & Koronakis, V. Arf6 coordinates actin assembly through the WAVE complex, a mechanism usurped by Salmonella to invade host cells. Proceedings of the National Academy of Sciences of the United States of America 110, 16880–16885 (2013).

12. Montagnac, G. et al. ARF6 Interacts with JIP4 to control a motor switch mechanism regulating endosome traffic in cytokinesis. Current biology : CB 19, 184–195 (2009).

13. Moravec, R., Conger, K.K., D’Souza, R., Allison, A.B. & Casanova, J.E. BRAG2/GEP100/IQSec1 interacts with clathrin and regulates alpha5beta1 integrin endocytosis through activation of ADP ribosylation factor 5 (Arf5). The Journal of biological chemistry 287, 31138–31147 (2012).

14. Rainero, E. et al. Ligand-Occupied Integrin Internalization Links Nutrient Signaling to Invasive Migration. Cell reports 10, 398–413 (2015).

15. Santy, L.C., Ravichandran, K.S. & Casanova, J.E. The DOCK180/Elmo complex couples ARNO-mediated Arf6 activation to the downstream activation of Rac1. Current biology : CB 15, 1749–1754 (2005).

16. Stamnes, M.A. & Rothman, J.E. The binding of AP-1 clathrin adaptor particles to Golgi membranes requires ADP-ribosylation factor, a small GTP-binding protein. Cell 73, 999–1005 (1993).

17. Donaldson, J.G. & Jackson, C.L. ARF family G proteins and their regulators: roles in membrane transport, development and disease. Nature reviews. Molecular cell biology 12, 362–375 (2011).

18. Hashimoto, A. et al. GEP100-Arf6-AMAP1-cortactin pathway frequently used in cancer invasion is activated by VEGFR2 to promote angiogenesis. PloS one 6, e23359 (2011).

19. Hashimoto, S. et al. Lysophosphatidic acid activates Arf6 to promote the mesenchymal malignancy of renal cancer. Nature communications 7, 10656 (2016).

20. Kinoshita, R. et al. Co-overexpression of GEP100 and AMAP1 proteins correlates with rapid local recurrence after breast conservative therapy. PloS one 8, e76791 (2013).

21. Morishige, M. et al. GEP100 links epidermal growth factor receptor signalling to Arf6 activation to induce breast cancer invasion. Nature cell biology 10, 85–92 (2008).

22. Hashimoto, S. et al. Targeting AMAP1 and cortactin binding bearing an atypical src homology 3/proline interface for prevention of breast cancer invasion and metastasis. Proceedings of the National Academy of Sciences of the United States of America 103, 7036–7041 (2006).

23. Grossmann, A.H. et al. The small GTPase ARF6 stimulates beta-catenin transcriptional activity during WNT5A-mediated melanoma invasion and metastasis. Science signaling 6, ra14 (2013).

24. Marchesin, V. et al. ARF6-JIP3/4 regulate endosomal tubules for MT1-MMP exocytosis in cancer invasion. The Journal of cell biology 211, 339–358 (2015).

25. Marchesin, V., Montagnac, G. & Chavrier, P. ARF6 promotes the formation of Rac1 and WAVE-dependent ventral F-actin rosettes in breast cancer cells in response to epidermal growth factor. PloS one 10, e0121747 (2015).

26. Muralidharan-Chari, V. et al. ADP-ribosylation factor 6 regulates tumorigenic and invasive properties in vivo. Cancer research 69, 2201–2209 (2009).

27. Zhang, Q. et al. Small-molecule synergist of the Wnt/beta-catenin signaling pathway. Proceedings of the National Academy of Sciences of the United States of America 104, 7444–7448 (2007).

28. Singh, M.K. et al. Structure-activity relationship studies of QS11, a small molecule Wnt synergistic agonist. Bioorganic & medicinal chemistry letters 25, 4838–4842 (2015).

29. Yoo, J.H. et al. ARF6 Is an Actionable Node that Orchestrates Oncogenic GNAQ Signaling in Uveal Melanoma. Cancer cell 29, 889–904 (2016).

30. Bello, D., Webber, M.M., Kleinman, H.K., Wartinger, D.D. & Rhim, J.S. Androgen responsive adult human prostatic epithelial cell lines immortalized by human papillomavirus 18. Carcinogenesis 18, 1215–1223 (1997).

31. Kaighn, M.E., Narayan, K.S., Ohnuki, Y., Lechner, J.F. & Jones, L.W. Establishment and characterization of a human prostatic carcinoma cell line (PC-3). Investigative urology 17, 16–23 (1979).

32. Someya, A. et al. ARF-GEP(100), a guanine nucleotide-exchange protein for ADP-ribosylation factor 6. Proceedings of the National Academy of Sciences of the United States of America 98, 2413–2418 (2001).

33. Dunphy, J.L., Ye, K. & Casanova, J.E. Nuclear functions of the Arf guanine nucleotide exchange factor BRAG2. Traffic 8, 661–672 (2007).

34. D’Souza, R.S. et al. Calcium-stimulated disassembly of focal adhesions mediated by an ORP3/IQSec1 complex. Elife 9 (2020).

35. Manavski, Y. et al. Brag2 differentially regulates beta1- and beta3-integrin-dependent adhesion in endothelial cells and is involved in developmental and pathological angiogenesis. Basic research in cardiology 109, 404 (2014).

36. Santy, L.C. Characterization of a fast cycling ADP-ribosylation factor 6 mutant. The Journal of biological chemistry 277, 40185–40188 (2002).

37. Beemiller, P., Hoppe, A.D. & Swanson, J.A. A phosphatidylinositol-3-kinase-dependent signal transition regulates ARF1 and ARF6 during Fcgamma receptor-mediated phagocytosis. PLoS biology 4, e162 (2006).

38. Hafner, M. et al. Inhibition of cytohesins by SecinH3 leads to hepatic insulin resistance. Nature 444, 941–944 (2006).

39. Benabdi, S. et al. Family-wide Analysis of the Inhibition of Arf Guanine Nucleotide Exchange Factors with Small Molecules: Evidence of Unique Inhibitory Profiles. Biochemistry 56, 5125–5133 (2017).

40. Lu, Z.X. et al. Transcriptome-wide landscape of pre-mRNA alternative splicing associated with metastatic colonization. Molecular cancer research : MCR 13, 305–318 (2015).

41. Roca, H. et al. Transcription factors OVOL1 and OVOL2 induce the mesenchymal to epithelial transition in human cancer. PloS one 8, e76773 (2013).

42. Lillis, A.P., Mikhailenko, I. & Strickland, D.K. Beyond endocytosis: LRP function in cell migration, proliferation and vascular permeability. Journal of thrombosis and haemostasis : JTH 3, 1884–1893 (2005).

43. Herz, J. et al. Surface location and high affinity for calcium of a 500-kd liver membrane protein closely related to the LDL-receptor suggest a physiological role as lipoprotein receptor. The EMBO journal 7, 4119–4127 (1988).

44. Li, H. et al. Targeting of mTORC2 prevents cell migration and promotes apoptosis in breast cancer. Breast cancer research and treatment 134, 1057–1066 (2012).

45. Zhang, F. et al. mTOR complex component Rictor interacts with PKCzeta and regulates cancer cell metastasis. Cancer research 70, 9360–9370 (2010).

46. Liu, P. et al. Sin1 phosphorylation impairs mTORC2 complex integrity and inhibits downstream Akt signalling to suppress tumorigenesis. Nature cell biology 15, 1340–1350 (2013).

47. Jian, X. et al. Autoinhibition of Arf GTPase-activating protein activity by the BAR domain in ASAP1. The Journal of biological chemistry 284, 1652–1663 (2009).

48. Lewis, S.M., Poon, P.P., Singer, R.A., Johnston, G.C. & Spang, A. The ArfGAP Glo3 is required for the generation of COPI vesicles. Molecular biology of the cell 15, 4064–4072 (2004).

49. Zhang, C.J., Cavenagh, M.M. & Kahn, R.A. A family of Arf effectors defined as suppressors of the loss of Arf function in the yeast Saccharomyces cerevisiae. The Journal of biological chemistry 273, 19792–19796 (1998).

50. Theret, L. et al. Identification of LRP-1 as an endocytosis and recycling receptor for beta1-integrin in thyroid cancer cells. Oncotarget 8, 78614–78632 (2017).

51. Kang, H.S. et al. LRP1-dependent pepsin clearance induced by 2’-hydroxycinnamaldehyde attenuates breast cancer cell invasion. The international journal of biochemistry & cell biology 53, 15–23 (2014).

52. Stephens, L. et al. Protein kinase B kinases that mediate phosphatidylinositol 3,4,5-trisphosphate-dependent activation of protein kinase B. Science 279, 710–714 (1998).

53. Sarbassov, D.D., Guertin, D.A., Ali, S.M. & Sabatini, D.M. Phosphorylation and regulation of Akt/PKB by the rictor-mTOR complex. Science 307, 1098–1101 (2005).

54. Brown, H.A., Gutowski, S., Moomaw, C.R., Slaughter, C. & Sternweis, P.C. ADP-ribosylation factor, a small GTP-dependent regulatory protein, stimulates phospholipase D activity. Cell 75, 1137–1144 (1993).

55. Cockcroft, S. et al. Phospholipase D: a downstream effector of ARF in granulocytes. Science 263, 523–526 (1994).

56. Honda, A. et al. Phosphatidylinositol 4-phosphate 5-kinase alpha is a downstream effector of the small G protein ARF6 in membrane ruffle formation. Cell 99, 521–532 (1999).

57. Yoo, J.H. et al. The small GTPase ARF6 activates PI3K in melanoma to induce a pro-metastatic state. Cancer research (2019).

58. Tsai, M.T. et al. Regulation of HGF-induced hepatocyte proliferation by the small GTPase Arf6 through the PIP2-producing enzyme PIP5K1A. Scientific reports 7, 9438 (2017).

59. Morran, D.C. et al. Targeting mTOR dependency in pancreatic cancer. Gut 63, 1481–1489 (2014).

60. Luo, L. et al. TLR Crosstalk Activates LRP1 to Recruit Rab8a and PI3Kgamma for Suppression of Inflammatory Responses. Cell reports 24, 3033–3044 (2018).

61. Volpicelli-Daley, L.A., Li, Y., Zhang, C.J. & Kahn, R.A. Isoform-selective effects of the depletion of ADP-ribosylation factors 1-5 on membrane traffic. Molecular biology of the cell 16, 4495–4508 (2005).

62. Zhu, W. et al. Small GTPase ARF6 controls VEGFR2 trafficking and signaling in diabetic retinopathy. The Journal of clinical investigation 127, 4569–4582 (2017).

63. Ratcliffe, C.D.H. et al. HGF-induced migration depends on the PI(3,4,5)P3-binding microexon-spliced variant of the Arf6 exchange factor cytohesin-1. The Journal of cell biology 218, 285–298 (2019).

64. Klarlund, J.K., Tsiaras, W., Holik, J.J., Chawla, A. & Czech, M.P. Distinct polyphosphoinositide binding selectivities for pleckstrin homology domains of GRP1-like proteins based on diglycine versus triglycine motifs. The Journal of biological chemistry 275, 32816–32821 (2000).

65. Ogasawara, M. et al. Similarities in function and gene structure of cytohesin-4 and cytohesin-1, guanine nucleotide-exchange proteins for ADP-ribosylation factors. The Journal of biological chemistry 275, 3221–3230 (2000).

66. Roman-Fernandez, A. et al. The phospholipid PI(3,4)P2 is an apical identity determinant. Nature communications 9, 5041 (2018).

67. Somers, K.D. et al. Orthotopic treatment model of prostate cancer and metastasis in the immunocompetent mouse: efficacy of flt3 ligand immunotherapy. International journal of cancer 107, 773–780 (2003).

68. Hubner, N.C. et al. Quantitative proteomics combined with BAC TransgeneOmics reveals in vivo protein interactions. The Journal of cell biology 189, 739–754 (2010).

69. Cox, J. & Mann, M. MaxQuant enables high peptide identification rates, individualized p.p.b.-range mass accuracies and proteome-wide protein quantification. Nature biotechnology 26, 1367–1372 (2008).

70. Cox, J. et al. Andromeda: a peptide search engine integrated into the MaxQuant environment. Journal of proteome research 10, 1794–1805 (2011).

71. UniProt, C. The Universal Protein Resource (UniProt) in 2010. Nucleic acids research 38, D142–148 (2010).

72. Cox, J. et al. Accurate proteome-wide label-free quantification by delayed normalization and maximal peptide ratio extraction, termed MaxLFQ. Molecular & cellular proteomics : MCP 13, 2513–2526 (2014).

73. Perez-Riverol, Y. et al. The PRIDE database and related tools and resources in 2019: improving support for quantification data. Nucleic acids research 47, D442–D450 (2019).

74. Cerami, E. et al. The cBio cancer genomics portal: an open platform for exploring multidimensional cancer genomics data. Cancer Discov 2, 401–404 (2012).

75. Gao, J. et al. Integrative analysis of complex cancer genomics and clinical profiles using the cBioPortal. Science signaling 6, pl1 (2013).

76. Sun, W. et al. TSVdb: a web-tool for TCGA splicing variants analysis. BMC Genomics 19, 405 (2018).

