## Supplementary Table S1 for "Spatial enrichment of phosphoinositide metabolism is a molecular switch to promote metastasis"

| Our nomenclature | UCSC Genome Browser | NCBI Nucleotide RefSeq | NCBI Nucleotide ID | NCBI Protein ID | Alternate IDs in literature | Additional IDs | Literature References |
|---|---|---|---|---|---|---|---|
| ***IQSEC1 V1*** | uc011auw.2 | NM_001134382.3 | IQSEC1a (Variant 1) | NP_001127854.1 | BRAG2c | | |
| ***IQSEC1 V2*** | uc003bxt.3 | NM_014869.8 | IQSEC1b (Variant 2) | NP_055684.3 | BRAG2b | | PMID: 16461286 |
| ***IQSEC1 V3*** | | AB018306.1 | | BAA34483.2 | BRAG2a/GEP100 | KIAA0763 | PMID: 16461286 |
| ***IQSEC1 V4*** | uc003bxu.4 | NM_001330619.3 | IQSEC1c (Variant 3) | NP_001317548.1 | BRAG2d | | |
