## Supplementary Table S2 for "Spatial enrichment of phosphoinositide metabolism is a molecular switch to promote metastasis"

| row ID | isoform_uc003bxt_IsoformRatio_ControlDivided_Log2 | isoform_uc003bxu_IsoformRatio_ControlDivided_Log2 | isoform_uc011auw_IsoformRatio_ControlDivided_Log2 | V2-V1 ratio_ControlDivided_Log2 | gene_IQSEC1_ControlDivided_Log2 | Unique count(PatientID)_NormalSamples | Unique count(PatientID)_TumourSamples |
|---|---|---|---|---|---|---|---|
| BLCA | -0.045456415 | 0.209611274 | -0.903577952 | 0.019021341 | -0.607159218 | 19 | 408 |
| BRCA | 0.122631432 | 0.034544488 | -0.660839226 | 0.193657029 | -0.702844225 | 112 | 1093 |
| CESC | -0.521267714 | 0.487147343 | -0.824827712 | -0.446615781 | -1.314178386 | 3 | 304 |
| CHOL | -0.41318462 | 0.146758172 | 0.388284839 | -0.431482945 | -0.861457875 | 9 | 36 |
| COAD | 0.399549077 | 0.002879936 | -0.60584768 | 0.481819533 | -0.123054277 | 41 | 285 |
| ESCA | -0.350112396 | 0.154569678 | -0.253812333 | -0.347971136 | -0.897886497 | 11 | 184 |
| KICH | -0.085043453 | 0.158128944 | -0.46683997 | -0.014710706 | 0.73155292 | 25 | 66 |
| KIRC | 0.029167757 | -0.068937133 | -0.173623068 | 0.044274408 | -0.154391013 | 72 | 533 |
| LIHC | -0.138656548 | -0.177742205 | 0.175692343 | -0.180191926 | 0.389870155 | 50 | 371 |
| LUAD | -5.08151E-05 | 0.010617902 | -0.357559901 | 0.028067778 | -0.928837934 | 59 | 515 |
| LUSC | 0.00993884 | 0.046609151 | -0.385405262 | 0.055411452 | -1.536205495 | 51 | 501 |
| PAAD | 0.096973991 | -0.199250455 | 0.05334858 | 0.065052227 | -0.739011779 | 4 | 178 |
| PCPG | 0.209482449 | -1.080290529 | 0.969397455 | -0.013182406 | -0.804612344 | 3 | 179 |
| PRAD | 0.093651861 | 0.060694432 | -0.954155063 | 0.186195283 | 0.152708894 | 52 | 497 |
| READ | 0.328489267 | 0.031837489 | -0.711635262 | 0.425494546 | -0.109078638 | 10 | 94 |
| SARC | -0.676832522 | 0.092489326 | 0.657979532 | -0.868265759 | 0.343523911 | 2 | 259 |
| STAD | -0.057266156 | 0.002486747 | -0.19169818 | -0.030493111 | -0.250600462 | 35 | 415 |
| THCA | 0.18063912 | -0.158931094 | -0.188643694 | 0.142381635 | -0.157754297 | 59 | 501 |
| THYM | 0.830397892 | -0.65514746 | -0.869303471 | 0.936274521 | -0.403747177 | 2 | 120 |
| UCEC | 0.123325986 | 0.164771241 | -1.083334919 | 0.284650082 | -1.004096626 | 24 | 177 |
| **Total** | | | | | | **643** | **6716** |
