## Supplementary Table S3 for "Spatial enrichment of phosphoinositide metabolism is a molecular switch to promote metastasis"

| TumorType | Unique count(SampleID) |
|---|---|
| ACC | 46 |
| BLCA | 344 |
| BRCA | 892 |
| CESC | 173 |
| CHOL | 30 |
| CORE | 498 |
| DLBC | 33 |
| ESCA | 126 |
| GBM | 244 |
| HNSC | 212 |
| KICH | 63 |
| KIRC | 478 |
| KIRP | 217 |
| LGG | 435 |
| LIHC | 184 |
| LUAD | 365 |
| LUSC | 328 |
| MESO | 63 |
| OV | 436 |
| PAAD | 123 |
| PCPG | 82 |
| PRAD | 352 |
| SARC | 226 |
| SKCM | 356 |
| STAD | 392 |
| TGCT | 122 |
| THCA | 380 |
| THYM | 90 |
| UCEC | 440 |
| UCS | 48 |
| UVM | 12 |
| **Total** | 7790 |
